# PP2C phosphatase Pic14 negatively regulates tomato Pto/Prf-triggered immunity by inhibiting MAPK activation

**DOI:** 10.1101/2024.06.03.597203

**Authors:** Joydeep Chakraborty, Guy Sobol, Fan Xia, Ning Zhang, Gregory B. Martin, Guido Sessa

## Abstract

Type 2C protein phosphatases (PP2Cs) are emerging as important regulators of plant immune responses, although little is known about how they might impact nucleotide-binding, leucine-rich repeat (NLR)-triggered immunity (NTI). We discovered that expression of the PP2C-immunity associated candidate 14 gene (*Pic14*) is induced upon activation of the Pto/Prf-mediated NTI response in tomato. Pto/Prf recognize the effector AvrPto translocated into plant cells by the pathogen *Pseudomonas syringae* pv. *tomato* (*Pst*) and activate a MAPK cascade and other responses which together confer resistance to bacterial speck disease. *Pic14* encodes a PP2C with an N-terminal kinase-interacting motif (KIM) and a C-terminal phosphatase domain. Upon inoculation with *Pst*-AvrPto, Pto/Prf-expressing tomato plants with loss-of-function mutations in *Pic14* developed less speck disease, specifically in older leaves, compared to wild-type plants. Transient expression of Pic14 in leaves of *Nicotiana benthamiana* and tomato inhibited cell death typically induced by Pto/Prf and the MAPK cascade members M3Kα and Mkk2. The cell death-suppressing activity of Pic14 was dependent on the KIM and the catalytic phosphatase domain. Pic14 inhibited M3Kα- and Mkk2-mediated activation of immunity-associated MAPKs and Pic14 was shown to be an active phosphatase that physically interacts with and dephosphorylates Mkk2 in a KIM-dependent manner. Together, our results reveal Pic14 as an important negative regulator of Pto/Prf-triggered immunity by interacting with and dephosphorylating Mkk2.

**SIGNIFICANCE STATEMENT:** Plant intracellular immune receptors, typically nucleotide-binding, leucine-rich repeat proteins (NLRs) such as the tomato Prf protein activate NLR-triggered immunity (NTI) in response to specific pathogen virulence proteins. This paper reveals how a protein phosphatase interacts with and dephosphorylates a key signaling component acting downstream of Pto/Prf, likely to moderate negative effects of NTI on growth or other plant processes.

## INTRODUCTION

Plants have an innate immune system that uses intracellular receptors to detect virulence proteins, called effectors, that pathogens translocate into host cells during the infection process. These receptors are nucleotide-binding leucine-rich repeat (NLR) proteins and are categorized into subclasses based on their variable N-terminal domains: CC-NLRs (CNL proteins) with a coiled-coil (CC) domain, TIR-NLRs (TNL proteins) with a Toll and interleukin-1 receptor (TIR) domain, and RPW8-NLRs (RNL proteins) with a RPW8-type CC (CC_R_) domain (Huang et al., 2023). Generally, CNLs and TNLs are considered to function as ‘sensor’ NLRs, which detect effectors, and RNLs are considered as ‘helper’ NLRs that act in downstream events (Chia & Carella, 2023; Huang et al., 2023). In tomato and other solanaceous plants, <u>N</u>LR required for <u>c</u>ell death (Nrc), i.e., Nrc2 and Nrc3 proteins, are helper NLRs that function redundantly to regulate Pto/Prf-mediated immune activation (Wu et al., 2017; Zhang et al., 2024). Upon their activation, some NLRs form large oligomeric complexes called resistosomes through their variable N-terminal domains (Chia & Carella, 2023). NLR-triggered immunity (NTI) activates numerous cellular events that are accompanied by a cascade of signaling events culminating in host cell death, termed the hypersensitive response (HR), that limits the spread of pathogens beyond the infected tissues (Contreras et al., 2023; Huang et al., 2023).

The pathogen *Pseudomonas syringae* pv. *tomato* (*Pst*) DC3000 infects tomato plants and causes bacterial speck disease (Pedley & Martin, 2003). To enhance its virulence, *Pst* DC3000 translocates ∼36 effectors into the plant cell cytoplasm to suppress immune responses and activate processes benefiting the pathogen (Nomura et al., 2023; Roussin-Léveillée et al., 2022; Wei et al., 2015). Wild relatives of tomato evolved to recognize two of these effectors, AvrPto and AvrPtoB, using the Pto kinase protein (Martin, 2011; Martin et al., 1993). Upon *Pst* DC3000 attack, Pto binds to AvrPto and AvrPtoB, preventing these effectors from acting on their intended virulence targets, and subsequently forms a complex with the CNL protein Prf (Cheng et al., 2011; Dong et al., 2009; Kim et al., 2002; Mucyn et al., 2006, 2009; Ntoukakis et al., 2013; Pedley & Martin, 2003; Sheikh et al., 2023; Xiang et al., 2008). Activation of the Pto/Prf complex are known to initiate the mitogen-activated protein kinase (MAPK) signaling cascades (Oh et al., 2010; Pedley & Martin, 2004, 2005; del Pozo et al., 2004; Roberts et al., 2019; Sheikh et al., 2023; Zhang et al., 2024). MAPK cascades are used by all eukaryotes and include three protein kinases that act in a sequential manner to transduce a signal intracellularly. The first protein is a MAP kinase kinase kinase (MAPKKK), followed by a MAP kinase kinase (MAPKK), and finally a MAP kinase (MAPK). MAPK signaling cascades regulate diverse processes, including plant growth, development, and responses to abiotic and biotic stresses (Su et al., 2018; Sun & Zhang, 2022; M. Zhang & Zhang, 2022).

Numerous MAPK cascades that play a role in immune responses have been elucidated in Arabidopsis, rice, tobacco, and tomato (Sun & Zhang, 2022). MAPKs that play a role in Pto/Prf-mediated immune signaling include two MAPKKK genes (M3Kα and M3Kε), two MAPKK genes (Mkk1 and Mkk2), and two MAPK genes (Mpk2 and Mpk3). Several of these components were shown to activate each other, making up a M3Kα-Mkk2-Mpk2/Mpk3 sequential module that ultimately leads to the HR and disease resistance (Ekengren et al., 2003; Melech-Bonfil & Sessa, 2010; Pedley & Martin, 2004; del Pozo et al., 2004). This module has been shown to be regulated by M3Kα-interacting protein 1 (Mai1), a receptor-like cytoplasmic kinase, that acts with M3Kα to positively contribute to PCD activation (Roberts et al., 2019). In addition, three 14-3-3 proteins, Tft1, Tft3, and Tft7, have been identified as interacting partners of M3Kα, with Tft7 interacting with Mkk2 as well (Oh et al., 2010; Oh & Martin, 2011b; Sheikh et al., 2023). When 14-3-3 proteins bind their target protein they can act as phosphorylation sensors and modulate their client protein’s activity, resulting in altered protein-protein interactions, subcellular localization, or conformation of target proteins (Lozano-Durán & Robatzek, 2015). Recently, it has been demonstrated that Tft7 helps to bridge the M3Kα-Mkk2 association, while Tft1 and Tft3 contribute to dissociating M3Kα from the Pto/Prf complex, allowing activation of the downstream MAPK module (Sheikh et al., 2023).

Protein phosphorylation plays a key role in immune signaling transduction and can facilitate protein-protein interactions or induce conformational changes in signaling components (Kadota et al., 2019). However, to maintain a balance between growth and defense, deactivation of immune signaling is essential (Huot et al., 2014). This process involves highly dynamic dephosphorylation events of immune-related protein kinases by protein phosphatases, including protein phosphatase 2Cs (PP2Cs). PP2Cs are serine/threonine phosphatases that contain a conserved C-terminal catalytic domain and a diverse N-terminal domain responsible for their localization, protein-protein interactions, and auto-inhibition of their own C-terminal PP2C domain (Shi, 2009). Some PP2C have a kinase-interacting motif (KIM) that has been shown to function as a docking mechanism (Brock et al., 2010; Schweighofer et al., 2007).

Several Arabidopsis PP2Cs are regulators of abscisic acid (ABA) signaling and some accomplish this by interacting with MAPKs via their KIM and affecting MAPK phosphorylation status (Bhaskara et al., 2019). For example, PP2C5 and AP2C1 are ABA-responsive PP2Cs with an N-terminal KIM that promotes interaction with MAPKs (Brock et al., 2010; Schweighofer et al., 2007). A *pp2c5 ap2c1* double mutant exhibits higher levels of MAPK activation after ABA treatment compared to wild-type plants (Brock et al., 2010). Other Arabidopsis PP2Cs include, ABA insensitive 1 (ABI1) that reduces MPK6 phosphorylation, thus impacting 1-aminocyclopropane 1-carboxylate synthase (ACS) induced ethylene production (Ludwików et al., 2014), and Highly ABA-Induced 1 (HAI1), HAI2, and HAI3 which interact with MPK3 and MPK6 to suppress their activation (Mine et al., 2017).

PP2Cs also interact with MAPKs in the context of NTI, although fewer examples are known. In *Arabidopsis*, the expression of HAI1 is induced by coronatine produced by *Pst* DC3000, and HAI1 mediated dephosphorylation of MPK3 and MPK6 suppresses plant immunity against *Pst* DC3000. However, NTI triggered by plant recognition of the *Pst* AvrRpt2 effector by CNL RPS2 could significantly inhibit the coronatine-mediated HAI1 induction and thereby promote the activation of MPK3, MPK6 and disease resistance (Mine et al., 2017). Interestingly, a recent study showed that two PP2Cs in *Arabidopsis,* AP2C1 and PP2C5 could directly interact with MPK3 and MPK6 and attenuate MAPK activation induced by AvrRpt2 (Wang et al., 2023).

Through an analysis of transcriptomic data of tomato, we identified a subset of PP2C-encoding genes whose transcript abundance increases during various immune responses (Sobol et al., 2022); we now refer to these as <u>P</u>P2C immunity-associated <u>c</u>andidate (Pic) genes. Previously, we showed that Pic1 suppresses pattern-triggered immunity by dephosphorylating the Pti1b kinase (Giska & Martin, 2019; Schwizer et al., 2017) and that Pic3 and Pic12 negatively regulate tomato defense in flagellin independent manner (Xia et al., 2024). Here, we build on the transcriptomic data analysis by Sobol et al. (2022) which revealed that 25 out of the 97 PP2C-encoding genes in tomato show differential expression during NTI. One of these PP2Cs, Pic14, was chosen for study because it carries a KIM typically required for interaction with MAPKs (Brock et al., 2010; Schweighofer et al., 2007; Singh et al., 2018). We report evidence that Pic14 is an active phosphatase that relies on its KIM to target the M3Kα-Mkk2-Mpk2/Mpk3 module to negatively regulate the Pto/Prf pathway.

## RESULTS

### Tomato loss-of-function RG-pic14 mutants show enhanced activation of NTI responses

Our earlier RNA-Seq analysis of two *prf* tomato mutants (RG-prf3 and RG-prf19) revealed that transcript abundance of *Pic14*, which encodes a PP2C with a putative kinase-interacting motif (KIM), increased 3- to 6-fold upon Pto/Prf-induced NTI activation (Sobol et al., 2022; Pombo et al., 2014) (Table S1). To confirm this, we measured *Pic14* mRNA abundance in wild-type (RG-PtoR) tomato after inoculation with *Pst* DC3000 or DC3000Δ*avrPto*Δ*avrPtoB (*DC3000ΔΔ); the former strain activates Pto/Prf NTI while the latter does not. *Pic14* transcript abundance was increased about 3-fold at 6 h after inoculation with DC3000 compared to DC3000ΔΔ (Figure 1a).

**Figure 1.**
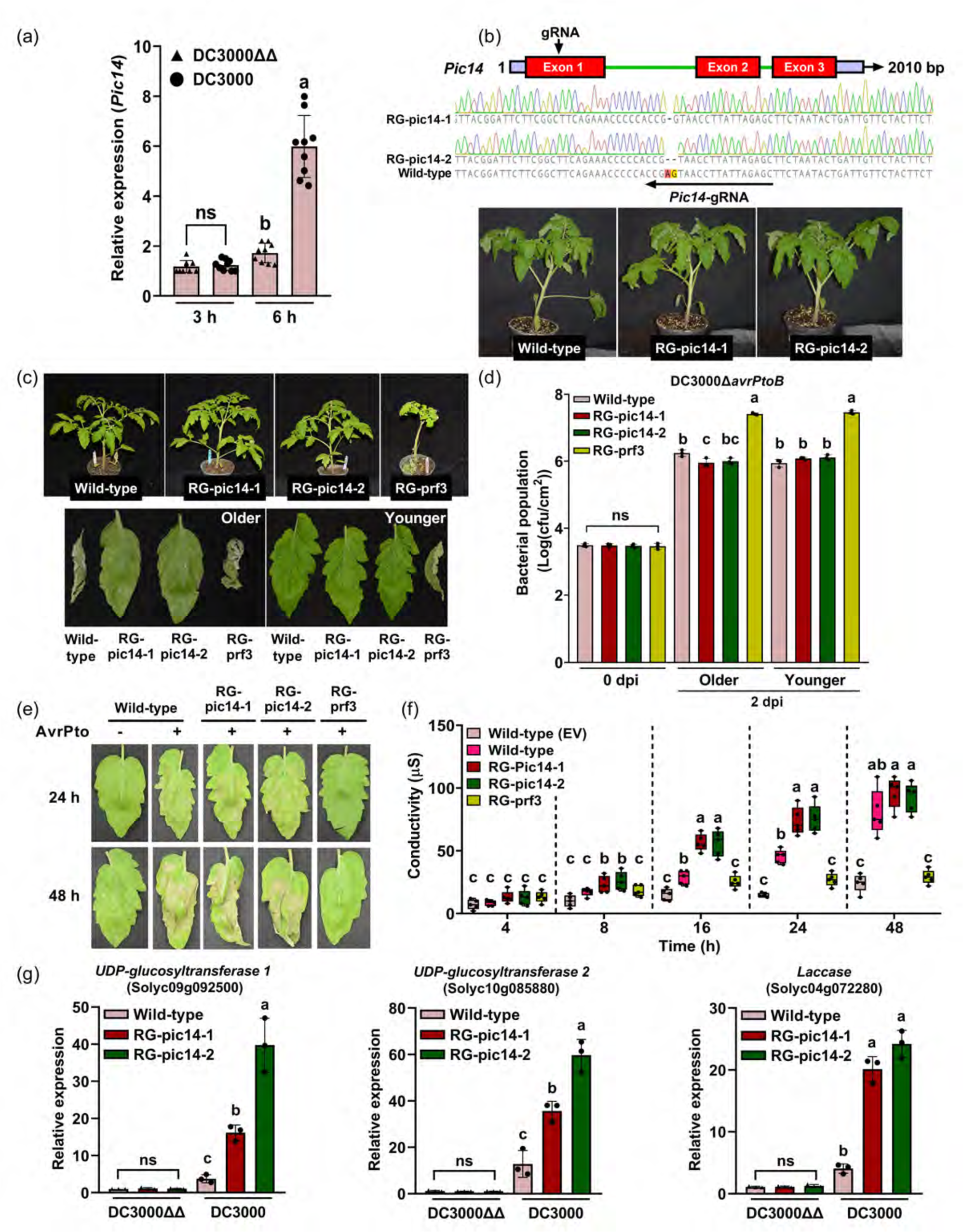
Tomato loss-of-function RG-pic14 mutants show enhanced activation of NTI responses. (a) Tomato *Pic14* mRNA accumulation is induced by the activation of Pto/Prf-mediated NTI. *Pic14* transcript abundance was measured by RT-qPCR following inoculation with *Pst* DC3000 and *Pst* DC3000Δ*avrPto*Δ*avrPtoB* (DC3000ΔΔ). Five-week-old plants of wild-type tomato cultivar Rio Grande-PtoR (RG-PtoR, which contains the *Pto* and *Prf* genes) were vacuum infiltrated with 5×10^6^ cfu mL^−1^ DC3000 or DC3000ΔΔ. cfu, colony-forming units. Bacteria-infiltrated tomato leaf samples were collected at 3 h and 6 h after vacuum infiltration. *SlEF1α* was used as the reference gene for RT-qPCR experiments. Error bars represent means ± standard deviation (SD); n = 3 plants with three technical repeats. The experiment was performed three times with similar results. (b) Top panel, *Pic14* gene structure, including exons (red) and introns (green) and the arrow points to the CRISPR/Cas9 guide-RNA (gRNA) target site. Purple bars indicate the 5’ and 3’ untranslated regions (UTRs) of *Pic14* gene. Middle panel, sequencing chromatogram of two missense mutant alleles generated by CRISPR/Cas9. The RG-pic14-1 line has a 1-bp deletion, and the RG-pic14-2 line has a 2-bp deletion. The deleted nucleotide bases are highlighted in orange (A) and yellow (G) in the sequence. The *Pic14*-gRNA sequence is indicated by a black arrow. Bottom panel, photographs of four-week-old wild-type (RG-PtoR) and RG-pic14 mutant plants. (c) Top panel, photos of wild-type (RG-PtoR), RG-pic14, and RG-prf3 plants that were vacuum infiltrated with 2×10^5^ cfu mL^−1^ DC3000Δ*avrPtoB*. Photographs were taken at 6 dpi. Bottom panel, older and younger leaf phenotypes at 6 dpi. ‘Older’ is the 1^st^ or 2^nd^ true leaf, and ‘younger’ refers to the 4^th^ or 5^th^ true leaf. The experiment was performed three times with similar results. (d) Measurement of DC3000Δ*avrPtoB* bacterial populations in wild-type (RG-PtoR), RG-pic14, and RG-prf3 plants. Four-week-old wild-type (RG-PtoR), RG-pic14, and RG-prf3 plants were vacuum infiltrated with 2×10^5^ cfu mL^−1^ DC3000Δ*avrPtoB*. Bacterial populations were measured at 3 hours (Day 0) and 2 days post-inoculation (dpi). Bars represent ± SD; n = 3. The experiment was performed three times with similar results. (e) AvrPto-induced cell death in wild-type (RG-PtoR), RG-pic14, and RG-prf3 plants. Older leaves of four-week-old wild-type (RG-PtoR), RG-pic14, and RG-prf3 plants were syringe infiltrated with *Agrobacterium tumefaciens* strain GV2260 containing pER8:EV (empty vector; -) or pER8:AvrPto (+) at OD_600_=0.3. Twenty-four hours later, the agroinfiltrated leaves were treated with 5 μM estradiol. Shown are representative leaf images at 24 h and 48 h after estradiol treatment. The experiment was performed twice with similar results. (f) Measurement of electrolyte leakage in wild-type (RG-PtoR), RG-pic14, and RG-prf3 older leaves in response to the estradiol-induced expression of pER8:AvrPto. Each data point is shown as the mean ± SD of n = 5 leaf discs. Similar results were obtained in two independent experiments. (g) NTI reporter gene expression in wild-type (RG-PtoR) and RG-pic14 mutants by RT-qPCR. Four-week-old wild-type (RG-PtoR) and RG-pic14 mutant tomato plants were vacuum infiltrated with 5×10^6^ cfu mL^−1^ DC3000ΔΔ or DC3000 strains, and leaf tissue was collected at 6 h post-infiltration. *SlEF1α* was used as the reference gene for RT-qPCR. Error bars represent means ± SD; n = 3 plants. The experiment was performed three times with similar results. In (a), (d), (f), and (g), different letters denote significant differences at the similar time point using one-way analysis of variance (ANOVA), followed by Tukey’s Honest Significant Difference (HSD) post hoc test (*P* < 0.05). ns, no significant difference.

Based on the increased abundance of *Pic14* transcripts during the Pto/Prf response, we hypothesized the Pic14 protein might play a role in regulating NTI and we therefore used CRISPR/Cas9 to generate mutations in the *Pic14* gene (Figure 1b). Two lines, RG-pic14-1 and RG-pic14-2, with each carrying a different homozygous mutation in *Pic14* were identified. RG-pic14-1 has a single base pair deletion in the first exon of the *Pic14* gene, resulting in the generation of a premature stop codon at the 87^th^ amino acid of the protein, while RG-pic14-2 has a two base pair deletion in the first exon of *Pic14*, causing an early stop codon at the 71^st^ amino acid (Figure S1). The overall growth, development, and morphology of the RG-pic14 mutants were indistinguishable from wild-type (RG-PtoR) plants (Figure 1b).

To assess the possible involvement of Pic14 in the Pto/Prf-triggered immunity, we used strain DC3000Δ*avrPtoB* strain which expresses AvrPto and not AvrPtoB; this strain has been shown previously to induce a weaker Pto/Prf NTI response than DC3000 which allows observation of milder impacts on NTI by host mutations (Zhang et al., 2024). Six days after inoculation with DC3000Δ*avrPtoB*, we observed no bacterial specks on either wild-type plants or the RG-pic14 mutants whereas RG-prf3 plants, as expected, developed severe disease symptoms (Figure 1c).

However, interestingly, the older leaves of wildtype plants wilted and abscised while no wilting of older leaves was observed in either of the RG-pic14 mutants (Figure 1c). Despite these differences in symptoms of the older leaves, we observed no statistically significant difference in bacterial populations in older or younger leaves of the RG-pic14 mutants as compared to wild-type (RG-PtoR) plants. RG-prf3 plants supported much larger bacterial populations than wildtype plants, as expected (Figure 1d). These results suggest that Pic14 might negatively regulates Pto/Prf-mediated NTI in older leaves and we therefore focused subsequent experiments on these leaves.

The recognition of AvrPto by Pto/Prf triggers HR cell death in tomato. To test if the loss-of-function mutations in Pic14 affect HR responses to AvrPto, we agroinfiltrated older leaves of wild-type RG-PtoR, RG-pic14, and RG-prf3 plants using an AvrPto construct under control of an estradiol-inducible system (Pedley & Martin, 2004). At 16 and 24 hours after estradiol treatment, RG-pic14 leaves exhibited increased cell death and electrolyte leakage compared to the wild-type (RG-PtoR) leaves (Figure 1e,f). Estradiol treatment resulted in no cell death in wild-type (RG-PtoR) leaves infiltrated with an empty vector (EV) control, while RG-prf3 leaves did not develop an HR after agroinfiltration with AvrPto as it lacks a functional Pto/Prf pathway.

We next assessed the transcript abundance of three NTI-specific genes, *UDPG1*, *UGPG2*, and *laccase*, that were shown previously to be induced 6 h after inoculation with DC3000 and not DC3000ΔΔ (Pombo et al., 2014). When compared to wild-type (RG-PtoR) plants, RG-pic14 mutants treated with DC3000 showed a significant increase in the relative abundance of *UDPG1*, *UGPG2*, and *laccase* transcripts at 6 h after inoculation (Figure 1g). These results further support the notion that Pic14 acts as a negative regulator of Pto/Prf-induced NTI responses in tomato.

### Pic14 suppresses host cell death associated with the CNLs Prf and Ptr1

Pic14 resides in clade B or clade E depending on the phylogenetic analysis of the tomato PP2Cs (Qiu et al., 2022; Sobol et al., 2022). Pic14 is a member of a small subgroup (of only 3 members) of tomato PP2C family phosphatases that has an N-terminal kinase-interacting motif (KIM) in addition to the C-terminal catalytic or PP2C domain (Figure 2a and Figure S2, Sobol et al., 2022). The KIM, consists of a consensus sequence of [K/R]_(3-4)_-X_(1-6)_-[L/I]-X-[L/I] (Brock et al., 2010; Schweighofer et al., 2007). Previously, PP2C5 and AP2C1, which both have the KIM domain were found to negatively affect MAPK activation during Arabidopsis immune responses and other processes (Ayatollahi et al., 2022; Brock et al., 2010; Diao et al., 2024; Schweighofer et al., 2007). The cell death mediated by Prf, a CNL, involves a MAPK cascade and we hypothesized therefore that Pic14 might affect this host response (Sheikh et al., 2023; Zhang et al., 2024). To investigate this issue, we first generated amino acid substitutions (K88A/R89Q) in the KIM to create the Pic14AQ variant and substitutions at D160N and D234N to generate the Pic14NN variant that was expected to abolish PP2C phosphatase activity (Figure 2a, Couto et al., 2016; Giska & Martin, 2019). A total of four plant expression constructs were made expressing YFP, Pic14, Pic14AQ, and Pic14NN each with an HA-epitope tag.

**Figure 2.**
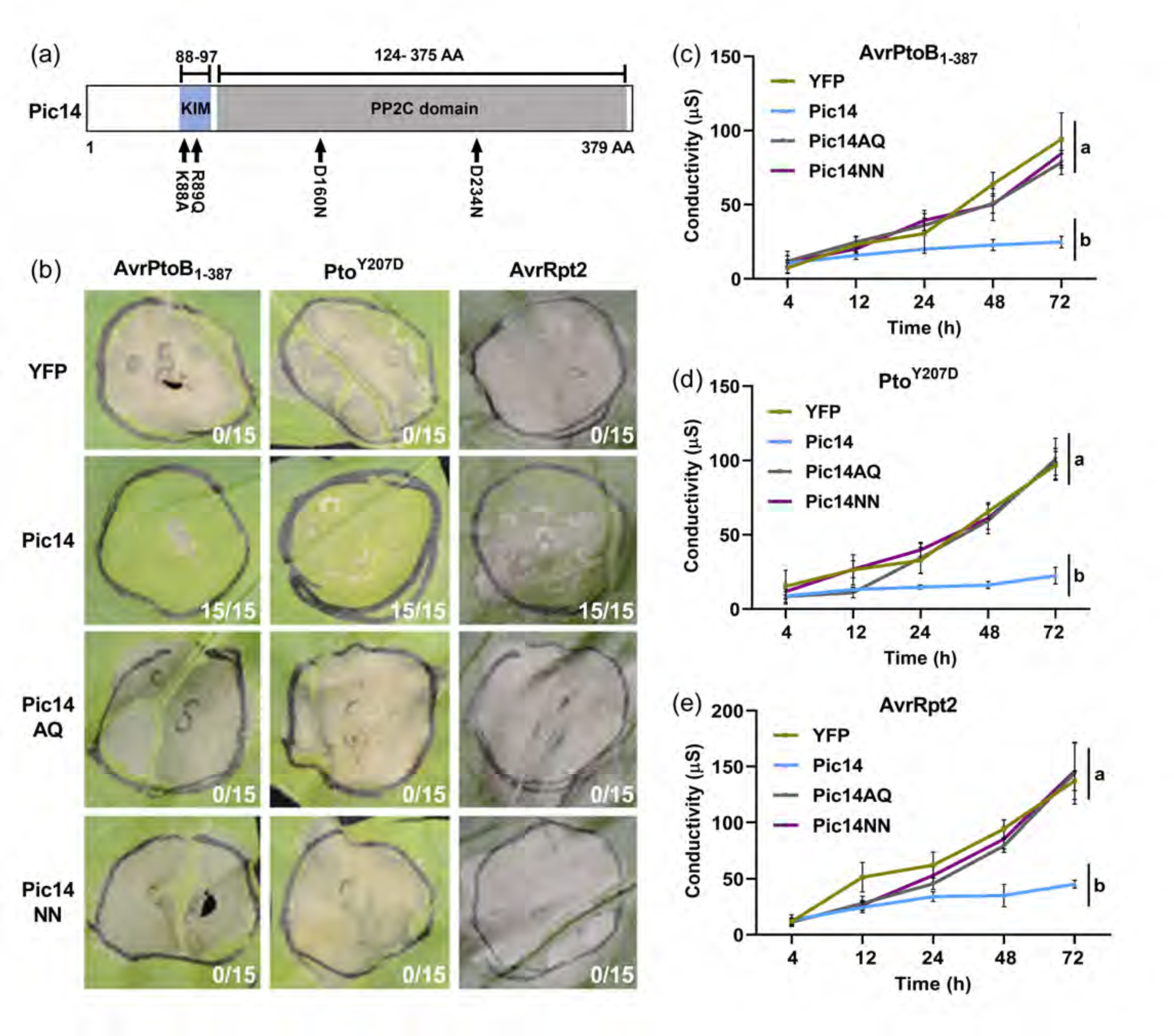
Pic14 suppresses cell death mediated by the Pto/Prf and Ptr1 pathways. (a) Diagram representing the protein domains of Pic14. Black arrows indicate the substituted amino acid (AA) residues from the kinase-interacting motif (KIM) and the PP2C domain. The numbers above indicate the span of KIM and PP2C domains within Pic14. Single-letter codes were used for substituted amino acids such as lysine (K), arginine (R), alanine (A), glutamine (Q), aspartic acid (D), and asparagine (N). (b) Pic14 inhibits cell death triggered by AvrPtoB_1-387_, Pto^Y207D^, and AvrRpt2. Leaves of five-week-old *Nicotiana benthamiana* plants were infiltrated with *A. tumefaciens* strains GV2260 containing YFP or Pic14, Pic14AQ, and Pic14NN variants at OD_600_=0.4. After 24 h, the infiltrated areas of the *N. benthamiana* leaves were subjected to another agroinfiltration with GV2260 strains carrying AvrPtoB_1-387_ or Pto^Y207D^ at OD_600_=0.4 or AvrRpt2 at OD_600_=0.2. The leaf images were taken 3 days post-infiltration with AvrPtoB_1-387_, AvrRpt2, or 7 days post-infiltration with Pto^Y207D^. Shown are the number of times cell death was observed to be suppressed by Pic14 or the variants (numerator) over the total number of agroinfiltrations performed (denominator). (c-e) Measurement of electrolyte leakage in *N. benthamiana* leaf discs co-expressing YFP or Pic14, Pic14AQ, and Pic14NN variants with cell death-inducing agents AvrPtoB_1-387_, Pto^Y207D^, and AvrRpt2. Each data point is shown as the mean ± SD of n = 5 leaves. Similar results were obtained in two independent experiments. Different letters indicate significant differences at a 72 h point using a one-way ANOVA followed by Tukey’s Honest Significant Difference (HSD) post hoc test (*P* < 0.05).

To assess the possible role of Pic14 in the Pto/Prf signaling pathway, *Agrobacterium tumefaciens* GV2260 strains carrying constructs of Pic14, the Pic14 variants or YFP were syringe-infiltrated into leaves of 5-week-old *N. benthamiana* plants and protein expression was confirmed (Figure S3). Then, 24 h later, the injected areas were agroinfiltrated with AvrPtoB_1-387_, a variant of AvrPtoB that elicits strong Pto/Prf-dependent NTI, or the autoactive variant protein, Pto^Y207D^ (Du et al., 2012; Oh et al., 2010). Pto^Y207D^ elicits Prf-dependent NTI responses and cell death in *N. benthamiana* leaves (Rathjen et al., 1999). We observed that Pic14 completely inhibited AvrPtoB_1-387_ and Pto^Y207D^-induced cell death in *N. benthamiana* leaves (Figure 2b). Areas agroinfiltrated with the Pic14AQ and Pic14NN variants or YFP did not affect this cell death. These observations, which were corroborated by electrolyte leakage assays (Figure 2c,d), indicated that Pic14 negatively regulates Pto/Prf-induced cell death and this activity requires both the KIM and the PP2C phosphatase domain.

We next investigated whether Pic14 plays a role in cell death mediated by two other NLRs, the CNL Ptr1 and the TNL Roq in response to the type III effectors AvrRpt2 or XopQ, respectively (Ahn et al., 2023; Mazo-Molina et al., 2019, 2020; Schultink et al., 2017). Leaves of *N. benthamiana,* which naturally expresses Ptr1, were agroinfiltrated with Pic14, the Pic14 variants or YFP and, 24 h later, agroinfiltrated with AvrRpt2. Pic14, but not Pic14AQ or Pic14NN, or YFP abolished cell death caused by AvrRpt2/Ptr1 and this was supported by electrolyte leakage assays (Figure 2b,e).

XopQ recognition by Roq results in weak cell death/chlorosis in *N. benthamiana* leaves which is monitored by the loss of the signal from a co-expressed GFP construct (Schultink et al., 2017a). We agroinfiltrated Pic14AQ and Pic14NN variants or YFP, and then 24 h later co-agroinfiltrated XopQ and GFP, or empty vector and GFP as a control (Figure S4). Expression of XopQ abolished GFP fluorescence and this was unaffected by co-expression of Pic14 or the Pic14 variants. GFP fluorescence was observed only in the empty vector control. Thus, Pic14 does not affect the Roq TNL-mediated cell death response. Collectively, these data indicate that Pic14 acts as a negative regulator of the immunity-associated cell death response induced by at least two CNL proteins.

### Host cell death associated with activation of M3Kα^KD^ and Mkk2 is suppressed by Pic14

The cell death induced by Pto/Prf NTI involves a MAPK cascade consisting of M3Ka, Mkk1/Mkk2, and MAP kinases, Mpk1, Mpk2, and Mpk3 (del Pozo et al., 2004; Oh & Martin, 2011b; Pedley & Martin, 2004). Expression of the kinase domain (KD) of M3Ka (M3Kα^KD^) alone in *N. benthamiana* leaves results in immunity-associated cell death (del Pozo et al., 2004; Sheikh et al., 2023). To determine whether Pic14 suppresses cell death induced by M3Kα^KD^, we agroinfiltrated Pic14, Pic14AQ, Pic14NN or YFP into *N. benthamiana* leaves and 24 h later agroinfiltrated M3Kα^KD^ under the control of an estradiol-inducible system (Figure S5a). Estradiol treatment caused M3Kα^KD^-triggered cell death in areas expressing Pic14AQ, Pic14NN, and YFP, but little or no cell death in Pic14-expressing regions (Figure 3a). These results were corroborated by an electrolyte leakage assay (Figure 3b).

**Figure 3.**
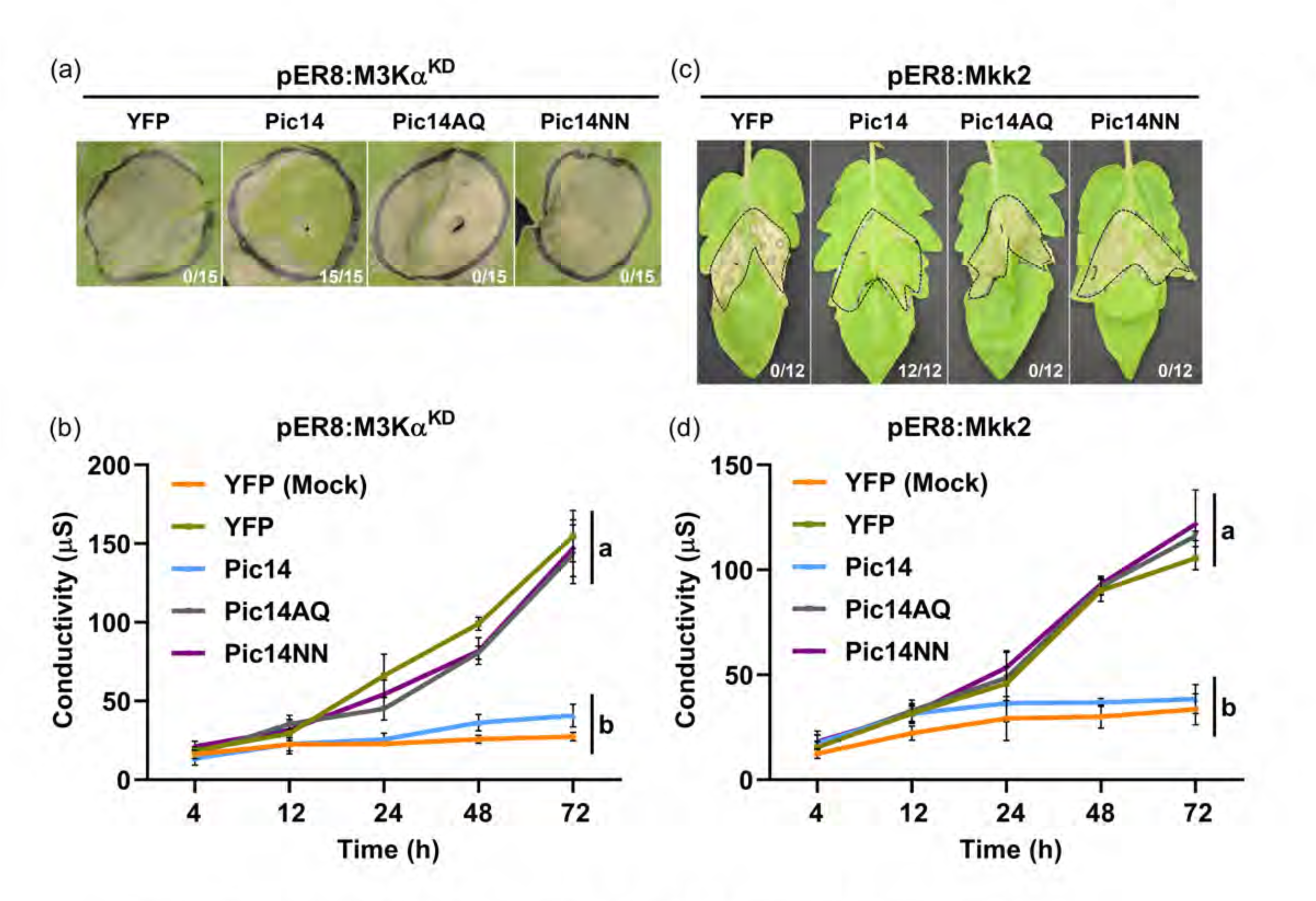
Pic14 suppresses M3Kα^KD^ and Mkk2-induced cell death in *N. benthamiana* and tomato leaves. (a) *A. tumefaciens* strains GV2260 carrying constructs of HA-epitope-tagged YFP or Pic14, Pic14AQ, and Pic14NN were infiltrated into *N. benthamiana* leaves at OD_600_=0.4. The next day, the infiltrated areas were infiltrated with GV2260 carrying pER8:M3Kα^KD^ at OD_600_=0.2. After 24 h post-infiltration, the abaxial surface of the leaves was sprayed with 5 μM estradiol. The image was taken 3 days after the estradiol treatment. Shown are the number of times cell death was observed to be suppressed by Pic14 or the variants (numerator) over the total number of agroinfiltrations performed (denominator). (b) Measurement of electrolyte leakage in the agroinfiltrated *N. benthamiana* leaves co-expressing YFP or Pic14 and its variants with M3Kα^KD^. (c) Pic14 inhibits cell death triggered by Mkk2 in wild-type (RG-PtoR) tomato leaves. Five-week-old tomato leaves were used for agroinfiltration with GV2260 strains containing constructs of YFP or Pic14, Pic14AQ, and Pic14NN variants at OD_600_=0.4. The next day, the agroinfiltrated leaves were further infiltrated with a GV2260 strain carrying pER8:Mkk2 at OD_600_=0.4. Twenty-four hours later, the infiltrated leaves were dipped into a solution containing 5 μM estradiol. Leaf images were captured 7 days post-estradiol treatment. Shown are the number of times cell death was observed to be suppressed by Pic14 or the variants (numerator) over the total number of agroinfiltrations performed (denominator). (d) Electrolyte leakage was measured in the agroinfiltrated tomato leaves co-expressing YFP, Pic14, Pic14AQ, and Pic14NN and Mkk2 at the indicated time points after estradiol treatment. In (b) and (d), a mock treatment was used as a negative control in the experiment. Each data point represents the mean ± SD of n = 5 infiltrated leaves. Different letters indicate significant differences at 72 h time point using a one-way ANOVA followed by Tukey’s Honest Significant Difference (HSD) post hoc test (*P* < 0.05). The experiment was repeated twice, with similar results.

Unlike M3Kα, Mkk2 does not cause immunity-associated cell death in *N. benthamiana* leaves but does so in tomato leaves (del Pozo et al., 2004; Pedley & Martin, 2004). We therefore agroinfiltrated Pic14, Pic14AQ, Pic14NN, or YFP into tomato leaves and 24 h later agroinfiltrated Mkk2 using the estradiol-inducible system. Mkk2-mediated cell death was suppressed in the Pic14-expressing areas, whereas cell death occurred in the Pic14AQ, Pic14NN, or YFP-expressing areas (Figure 3c). These results were corroborated by an electrolyte leakage assay (Figure 3d). Immunoblotting analysis confirmed the expression of M3Kα^KD^, Mkk2 in *N. benthamiana*, and YFP, Pic14, Pic14AQ, and Pic14NN expression in tomato leaves (Figure S5a,b). Together these results suggest that Pic14 negatively regulates Pto/Prf-mediated cell death by acting downstream of the NTI recognition complex and possibly directly on one or more components of the M3Ka-Mkk1/Mkk2-Mpk1/2/3 cascade.

### Pic14 interacts with Mkk2 in plant cells and the KIM is required for this interaction

Based on our observations above, we hypothesized that Pic14 might physically interact with a component of the M3Ka-Mkk1/Mkk2-Mpk1/2/3 cascade. Based on phylogenetic analysis, the most closely related proteins to Pic14 in other species appear to be *Zm*PP84 and *At*AP2C1 from maize and Arabidopsis, respectively (Figure S6a,b). *Zm*PP84 interacts with maize Mkk1 and *At*AP2C1 interacts with Mpk4 and Mpk6 in *Arabidopsis* (Guo et al., 2023; Schweighofer et al., 2007). We therefore focused on testing the possible interaction of Pic14 with Mkk1, Mkk2, Mpk1, Mpk2, and Mpk3 using a split-luciferase complementation assay (SLCA). Pic14, Pic14AQ, and Pic14NN were fused to the C-terminal half of the luciferase protein (C-LUC), and Mkk and Mpk proteins were fused to the N-terminal half of the luciferase protein (N-LUC). Using agroinfiltration, each C-LUC-Pic14 protein was co-expressed with each Mkk- and Mpk-N-LUC protein in *N. benthamiana* leaves (Figure 4a). As a negative control, C-LUC-Pic14 was co-expressed with the N-LUC empty vector (EV). After 48 hours, the agroinfiltrated leaves were collected to prepare leaf discs and incubated with luciferin substrate to measure the luminescence. We observed luminescence in the areas co-expressing Pic14 or Pic14NN with Mkk1, Mkk2 and Mpk1, however, the interaction between Pic14 or Pic14NN and Mkk2 was significantly stronger than with Mkk1 or Mpk1 (Figure 4a). The Pic14AQ protein did not interact with any of the Mkk or Mpk proteins indicating that the KIM domain is essential for interaction between Pic14 and Mkk1/Mkk2/Mpk1. A similar SLCA was performed on intact *N. benthamiana* leaves to allow comparisons on a single leaf. Again, the strongest luminescence resulted from the interaction between Pic14/Pic14NN and Mkk2 with a weaker interaction observable between Pic14/Pic14NN and Mkk1; co-expression of Pic14/Pic14NN with Mpk1 resulted in no observable luminescence (Figure 4b). Immunoblot analysis confirmed expression of each of the N- and C-LUC-tagged proteins in *N. benthamiana* leaves (Figure S7).

**Figure 4.**
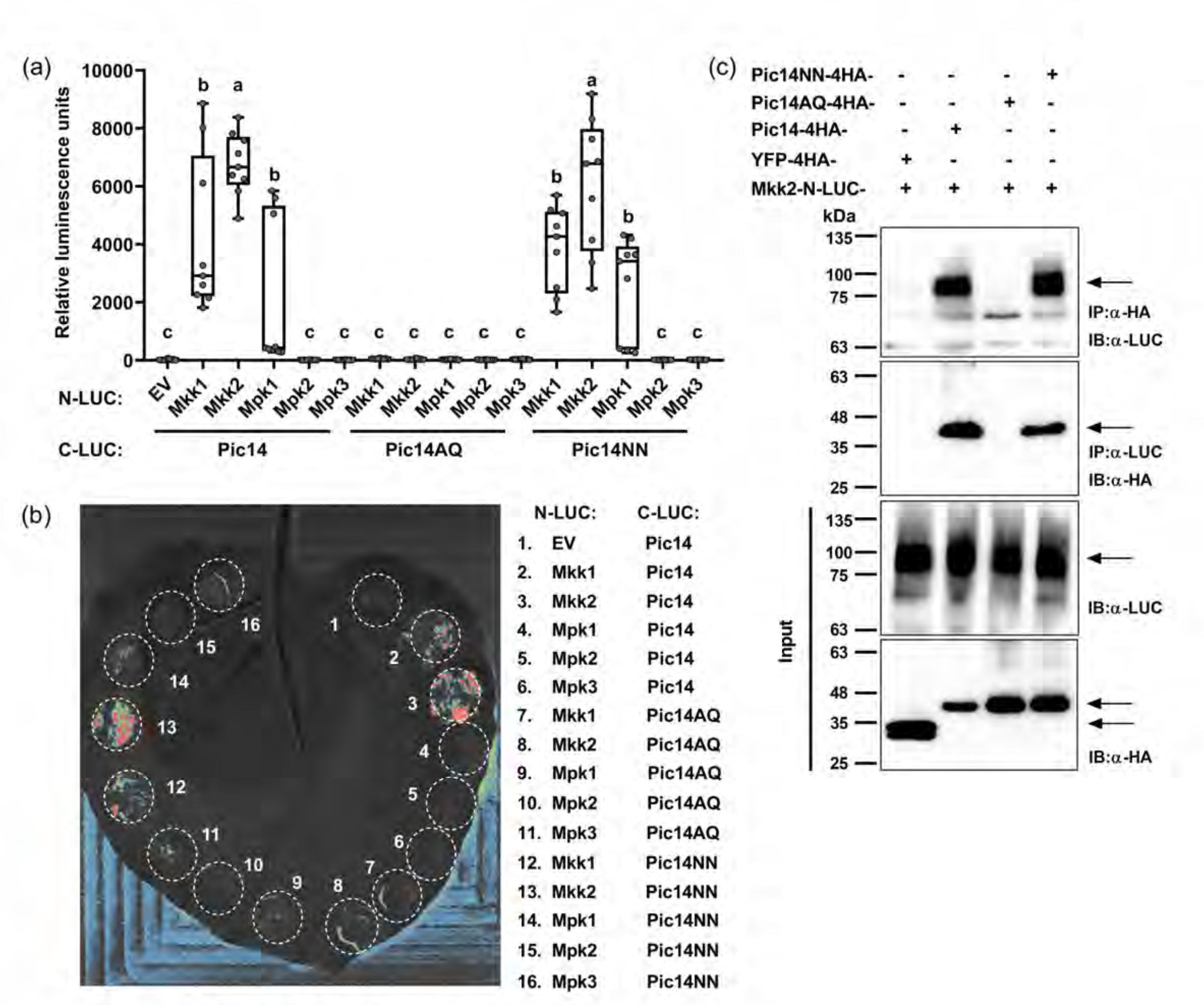
Pic14 interacts with Mkk2 in plant cells. (a) Quantification of the interaction between Pic14/Pic14AQ/Pic14NN and Mkk1/Mkk2/Mpk1/Mpk2/Mpk3, respectively, by split-luciferase complementation assay. *A. tumefaciens* GV2260 strains were used for co-expression of the C-LUC-Pic14/Pic14AQ/Pic14NN and N-LUC-empty vector (EV)/Mkk1/Mkk2/Mpk1/Mpk2/Mpk3 in leaves of five-week-old *N. benthamiana* plants. 48 hours after agroinfiltration, luciferase activity was measured by incubating the leaf discs with luciferin substrates. Luciferase activity was quantified as relative luminescence units. Data are means ± standard error (SE) of n = 9 leaf discs. Different letters indicate significant differences at a similar time point using a one-way ANOVA followed by Tukey’s Honest Significant Difference (HSD) post hoc test (*P* < 0.05). Similar results were obtained in three biological experiments. (b) A representative split-luciferase complementation assay image of an agroinfiltrated *N. benthamiana* leaf shows luminescence generated by superimposition of bright and dark field images. White circles indicate the infiltrated areas in the *N. benthamiana* leaf. The panel on the right side indicates the numbers that are displayed in the infiltration region to show the combinations of the constructs used for the co-expression of the proteins in *N. benthamiana*. The experiment was repeated two times with similar results. (c) Coimmunoprecipitation assay to determine association of Pic14 with Mkk2 in plant cells. *A. tumefaciens* GV2260 strains carrying 4HA-tagged YFP or Pic14, Pic14AQ, and Pic14NN variants were transiently co-expressed with Mkk2-N-LUC in leaves of five-week-old *N. benthamiana* plants. Following this, total leaf proteins were extracted and used for immunoprecipitation using anti-HA agarose beads and anti-LUC antibodies, respectively. The immunoblots were performed with anti-HA and anti-LUC antibodies. The (+) and (−) signs denote the presence or absence of the indicated proteins. Arrows indicate the specific protein bands after immunoblotting. This experiment was performed twice with similar results.

To further test the interaction between Pic14 and Mkk2, we performed *Agrobacterium*-mediated transient expression and co-immunoprecipitation (co-IP) of HA epitope-tagged Pic14, Pic14AQ, Pic14NN, and YFP with Mkk2-N-LUC in *N. benthamiana* leaves. In reciprocal co-IP experiments using either the anti-HA antibody or the anti-LUC antibody to precipitate the cognate proteins, we observed evidence of interaction of the Mkk2 protein with Pic14 and Pic14NN, but not with Pic14AQ (Figure 4c). Input samples showed the co-expression of the fusion proteins as detected by immunoblotting with anti-LUC and anti-HA antibodies. Together, these observations indicate that Pic14 uses its KIM to interact with Mkk2 and suggest that Mkk2 is a candidate for being an important substrate of Pic14.

### Pic14 inhibits MAPK phosphorylation associated with M3Kα^KD^ and Mkk2 activation in plant cells

It has been reported previously that overexpression of upstream signaling proteins such as M3Kα and Mkk2 in *N. benthamiana* and tomato can cause activation of downstream MAPKs (Gao et al., 2022; Pedley & Martin, 2004). To test whether Pic14 suppressed MAPK activation due to M3Kα or Mkk2 overexpression we agroinfiltrated Pic14, Pic14AQ, Pic14NN, or YFP into leaves of *N. benthamiana* and tomato, and 24 h later agroinfiltrated in the same areas estradiol-inducible M3Kα^KD^ and Mkk2. Upon estradiol treatment, MAPK activation was observed in leaf discs co-expressing M3Kα^KD^ or Mkk2 and YFP (Figure 5a,b). However, expression of Pic14 completely inhibited MAPK activation by M3Kα^KD^ and Mkk2 in *N. benthamiana* and tomato leaves, respectively (Figure 5a,b). Notably, co-expression of Pic14AQ and Pic14NN with M3Kα^KD^ and Mkk2 had no effect on MAPK activation. These results suggest that the Pic14 phosphatase activity and its KIM are required to inhibit MAPK activation, which in turn suppresses the HR in *N. benthamiana* and tomato leaves.

**Figure 5.**
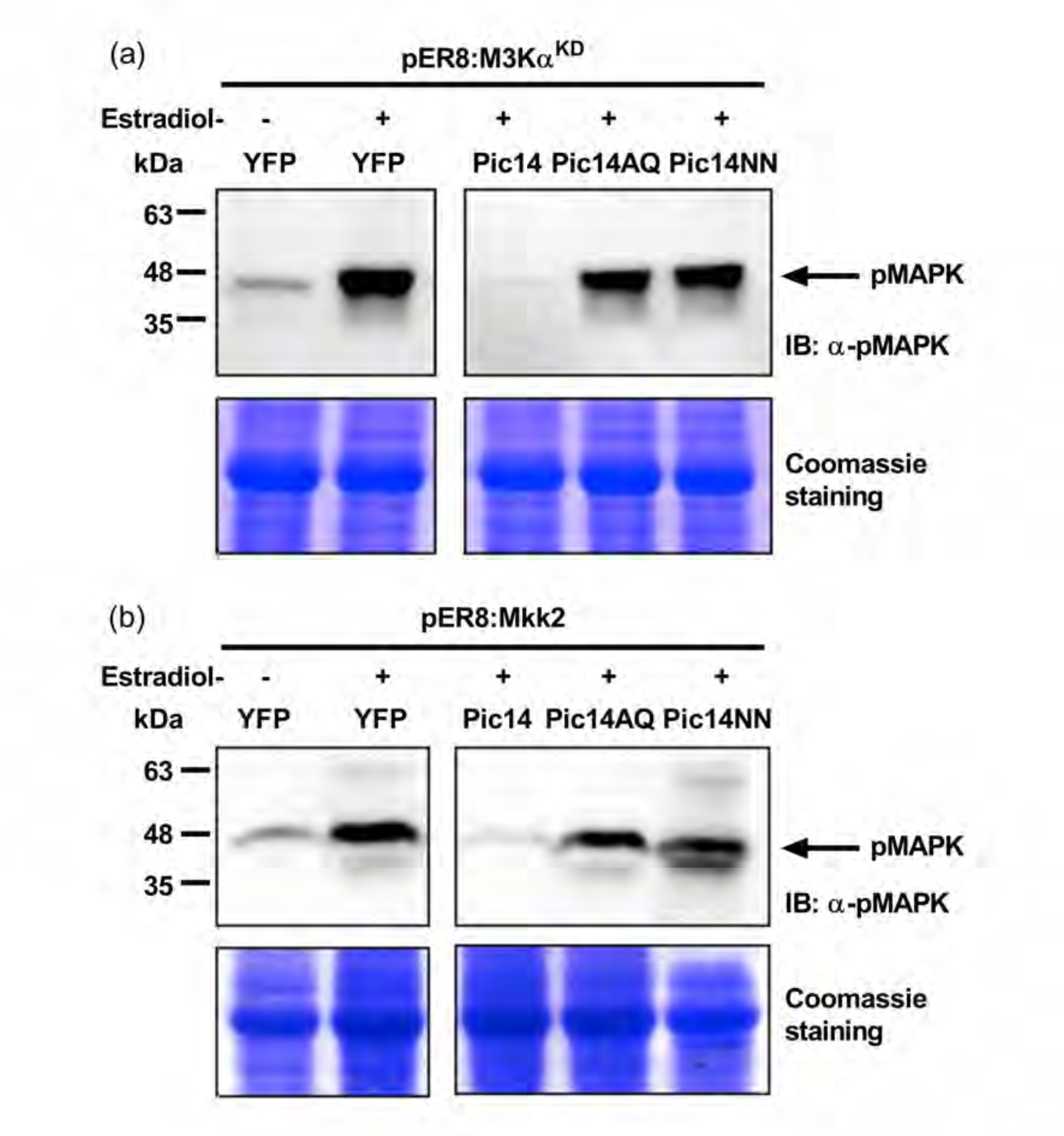
Pic14 suppresses MAPK activation induced by M3Kα^KD^ and Mkk2. (a) *Agrobacterium-*mediated transient expression of Pic14 inhibits MAPK phosphorylation induced by M3Kα^KD^ in *N. benthamiana* leaves. (b) Mkk2-induced MAPK activation in wild-type (RG-PtoR) tomato leaves is inhibited by Pic14. Five-week-old *N. benthamiana* or tomato leaves were first infiltrated with *A. tumefaciens* GV2260 strains carrying constructs of HA-tagged YFP or Pic14, Pic14AQ, and Pic14NN variants. The next day, infiltrated sites were syringe-infiltrated with pER8:M3Kα^KD^ or pER8:Mkk2. Following this, agroinfiltrated *N. benthamiana* and tomato leaves were treated with 5 μM estradiol. The leaf discs were harvested at 8 h after estradiol treatment. The (−) and (+) signs indicate mock treatment and estradiol-treated leaves, respectively. Total proteins were extracted from a pool of infiltrated leaf discs from three plants and used for immunoblotting by an anti-pMAPK antibody that binds to the phosphorylated MAPKs. Arrows indicate phosphorylated MAPK protein bands. Coomassie-stained gel serves as a loading control to show equal loading of protein. In (a) and (b), the images are derived from the same immunoblot with identical exposure times. Each experiment was repeated twice with similar results.

### Pic14 is an active phosphatase and dephosphorylates Mkk2 in a KIM-dependent manner

Based on its KIM-dependent association with Mkk2, we anticipated that Pic14, is an active phosphatase and likely dephosphorylates Mkk2. To test this, biochemical assays were used to assess the *in vitro* phosphatase activity of Pic14, expressed in *E. coli* as GST fusion proteins (Figure S8a). The recombinant proteins were incubated with a synthetic Ser/Thr phosphopeptide, and the release of inorganic phosphate was measured by a colorimetric assay. The result showed that GST-Pic14 and GST-Pic14AQ variants had significant phosphatase activity compared to the control GST protein, indicating the KIM domain of Pic14 is not required for binding the synthetic phosphopeptide (Figure 6a). GST-Pic14NN, however, exhibited no phosphatase activity, indicating that GST-Pic14NN is an inactive enzyme, which is consistent with earlier work (Couto et al., 2016; Giska & Martin, 2019).

**Figure 6.**
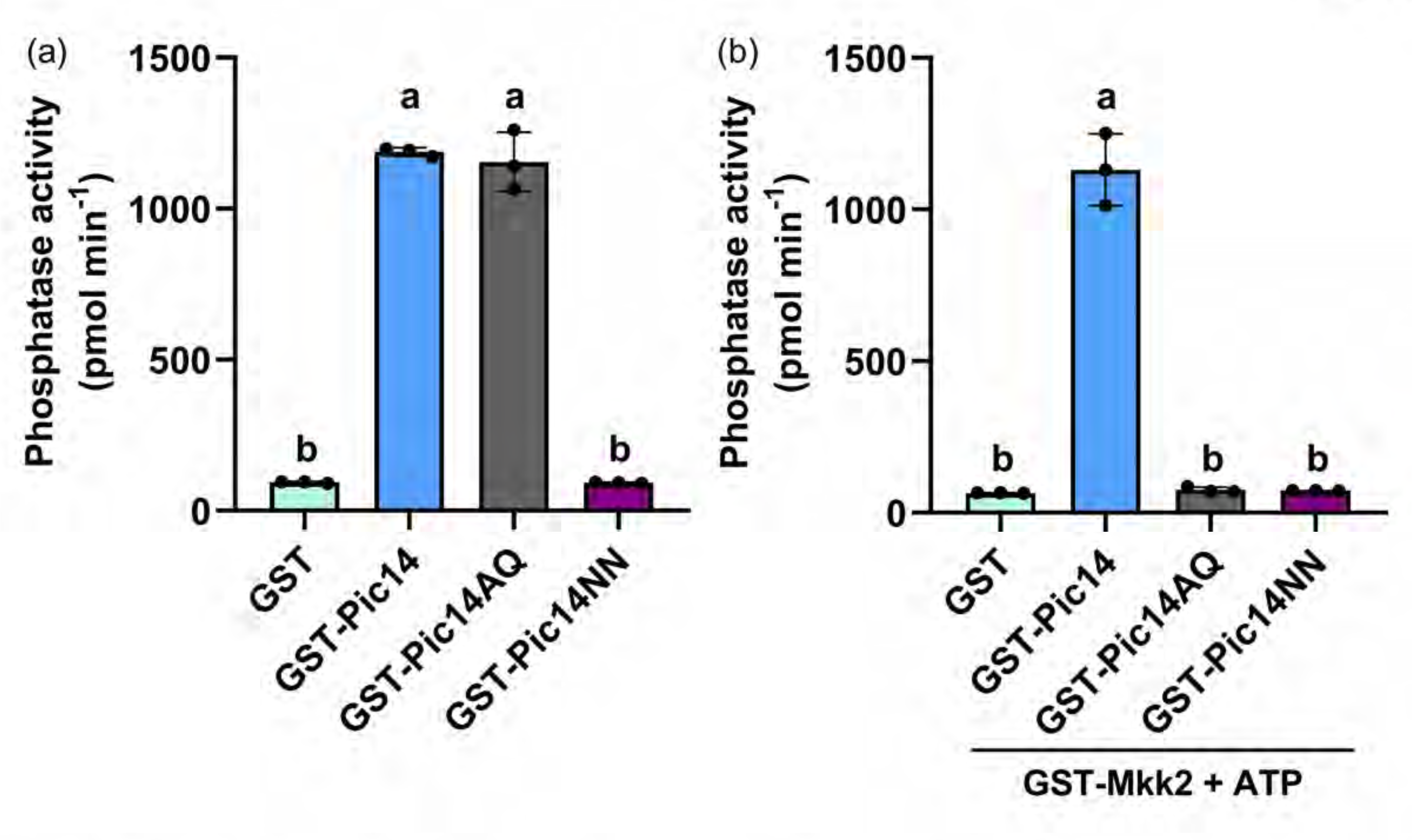
Pic14 is an active phosphatase that dephosphorylates Mkk2 in a KIM-dependent manner. (a) *In vitro* phosphatase activity of Pic14. The recombinant glutathione S-transferase (GST) fusion Pic14, Pic14AQ, and Pic14NN variants were purified from the Rosetta *Escherichia coli* strain, and the phosphatase activity was measured by a Ser/Thr phosphatase assay system (see Methods). GST alone was used as a negative control. (b) Mkk2 is dephosphorylated by Pic14 *in vitro*. GST-Mkk2 was purified from the Rosetta *E. coli* strain and subjected to *in vitro* phosphorylation in kinase assay buffer with 100 µM ATP for 30 min at 30°C. *In vitro* phosphorylated GST-Mkk2 protein was incubated with control GST or GST-Pic14, GST-Pic14AQ, and GST-Pic14NN variants, respectively, in a PP2C reaction buffer. The dephosphorylation activity of GST-Pic14 and its variants on the phosphorylated GST-Mkk2 protein was measured by the colorimetric method. In (a) and (b), error bars represent the means ± SD of n = 3 technical replicates. Three independent experiments were performed with similar results. Different letters denote significant differences in each data point using a one-way ANOVA followed by Tukey’s Honest Significant Difference (HSD) post hoc test (*P* < 0.05).

Next, we used the same Ser/Thr phosphatase assay to determine if Mkk2 could be dephosphorylated by GST-Pic14. To test this, we expressed and purified the Mkk2 protein as a GST fusion protein from *E. coli* (Figure S8b). *In vitro* phosphorylation of GST-Mkk2 proteins was carried out by incubating with cold ATP in a kinase assay buffer for thirty minutes. The reaction mixtures were incubated with control GST, GST-Pic14, GST-Pic14AQ, and GST-Pic14NN variants. The molybdate dye solution was added to the reaction and the OD was measured at 630 nm (Figure S8c). The dephosphorylation of recombinant Mkk2 protein was significantly increased when incubated with GST-Pic14 compared to the control GST protein, indicating that Mkk2 can be a substrate of Pic14 (Figure 6b). Mkk2 dephosphorylation was not detected with GST-Pic14AQ and GST-Pic14NN variants. Because GST-Pic14 did not show any phosphatase activity on GST-Mkk2 protein when incubated in a kinase assay buffer without the presence of ATP, indicating that the GST-Mkk2 protein is not phosphorylated in *E. coli* (Figure S8d,e). These results suggest that the KIM and PP2C domains are required for interaction Pic14 with Mkk2 and dephosphorylation of Mkk2, respectively.

## DISCUSSION

Protein phosphorylation serves as an important biochemical mechanism for plants and, in particular, acts as a switch to trigger and regulate immune responses (Kadota et al., 2019). Although much is known about the involvement of protein kinases in plant immunity, the role of protein phosphatases in regulating immune signaling remains less explored (Couto et al., 2016; Giska & Martin, 2019; Park et al., 2008; Schweighofer et al., 2007). Here we report on an active tomato protein PP2C phosphatase, Pic14, that negatively regulates Pto/Prf-mediated NTI by dephosphorylating a key component of an immunity-associated MAPK cascade. The initial indication for the involvement of Pic14 in NTI came from its increased transcript abundance upon Pto/Prf-induced NTI activation. Subsequent analysis of two independent CRISPR/Cas9-generated RG-pic14 mutants using bacterial inoculation assays revealed that Pic14 plays a negative regulatory role in Pto/Prf-induced NTI. By using the *Pst* DC3000Δ*avrPtoB* strain which allows observation of subtle effects on NTI, we detected increased resistance in older leaves in RG-pic14 mutants compared to wild-type plants, indicating an enhanced activation of immune responses in the absence of Pic14. This observation led to the discovery of the mechanism by which Pic14 inhibits Pto/Prf-triggered NTI through dephosphorylation of Mkk2, which likely inhibits subsequent Mpk phosphorylation. Here we discuss the leaf-age-related effects of Pic14 and its relation to other components of the Pto/Prf pathway, consider possible redundancy of Pic14 with a related PP2C, elaborate on the involvement of the KIM and phosphatase domain in the Pic14 mechanism of suppressing NTI, and speculate on why Pic14 acts differently in CNL- and TNL-mediated immunity.

Phenotypic evaluation of plants inoculated with *Pst* DC3000Δ*avrPtoB* demonstrated that RG-pic14 mutants are more resistant to bacterial infection in their older but not younger leaves. In a related study, we observed a similar phenotype in plants carrying mutations in two PP2Cs sharing sequence similarity, Pic3 and Pic12 (Xia et al., 2024). In that work, RNA-seq analysis revealed that older leaves of RG-pic3 had significantly more differentially expressed genes related to defense compared to younger leaves. Leaf age-dependent responses to *Pst* DC3000 have also been reported in Arabidopsis, supporting the notion of crosstalk between immune and developmental responses (Huot et al., 2014; Zipfel et al., 2004). Our additional experiments showing that AvrPto-induced cell death, electrolyte leakage, and transcript abundance of NTI-specific genes were all enhanced in RG-pic14 mutants further supports a role for Pic14 in the Pto/Prf-mediated NTI pathway. Other components of the Pto/Prf pathway such as Nrc2, Nrc3, Mai1, and Tft7 were characterized using similar assays, collectively supporting their involvement in the same pathway (Oh & Martin, 2011b; Roberts et al., 2019; Sheikh et al., 2023; Zhang et al., 2024).

In a related study we found that two closely related PP2Cs, Pic3 and Pic12, play a partially redundant role in non-NTI plant immune responses (Xia et al., 2024). Pic14 is also related to another PP2C, Pic6 (Solyc05g052520), with both PP2Cs residing in clade E (Qiu et al., 2022) or clade B (Sobol et al., 2022) depending on the phylogenetic analysis and sharing 55% amino acid sequence identity. Both genes are expressed in similar tomato tissues (Qiu et al., 2022). In common with Pic14, Pic6 also contains a KIM and C-terminal catalytic domain (Sobol et al., 2022). It is possible, therefore, that Pic6 and Pic14 play a partially redundant role in negatively regulating Pto/Prf-mediated NTI. Redundancy was also observed with another pair of PP2C proteins, the Arabidopsis AP2C1 and PP2C5, in the context of their effect on ABA-induced MAPK activation (Brock et al., 2010; Schweighofer et al., 2007). Upon ABA treatment, MAPK activation is higher in an *ap2C1/pp2C5* double mutant compared to the two single mutants (Brock et al., 2010). If Pic14 and Pic6 are partially redundant, it might explain why we observed relatively subtle increase in plant immunity in the RG-pic14 mutant lines when they were inoculated with DC3000Δ*avrPtoB*. A critical test of possible Pic6/Pic14 redundancy will require the future generation and characterization of RG-pic6 and RG-pic6/pic14 double mutants.

The N-terminal KIM and C-terminal PP2C domain are both essential for cell death suppression by Pic14. A KIM is also present in tomato MAPKKs and mediates their association with upstream and downstream components (Oh et al., 2013). However, proteins carrying both a KIM and a PP2C domain are unique to plants and not present in other eukaryotes (Fuchs et al., 2013; Schweighofer et al., 2007), suggesting an adaptation of some plant PP2Cs to enable modulation of phosphorylation status of MAPKs. The amino acid sequence of the Pic14 KIM is highly similar to the KIM present in other plant PP2Cs which are known to bind MAPKs in a KIM-dependent manner (Figure S2; Brock et al., 2010; Schweighofer et al., 2007), and Pic14 interacted with several MAPK cascade components including Mkk1, Mkk2, and Mpk1 in a KIM-dependent manner. We focused on the interaction of Pic14 with Mkk2 since it appeared to be the strongest and found, using *in vitro* phosphatase assays, that both the KIM and the PP2C domain are required for Pic14 dephosphorylation of Mkk2. Further studies will be required to determine whether Pic14 also dephosphorylates Mkk1 and Mpk1 despite their apparent weaker interaction. In the case that Mkk1 and Mpk1 are not substrates of Pic14 then the weak interactions might be due to the fact that a split luciferase complementation assay is proximity based, which could indicate a false positive interaction for a protein that is closely associated with the real interactor. In the future, it will be interesting to characterize interaction dynamics between Pic14 and its substrates before and after NTI activation. This will address the possibility that the interaction between Pic14 and its substrates is constitutive in order to suppress NTI and whether Pic14 is then released in response to an activation signal to allow NTI activation, similar to Arabidopsis PP2C38 (Couto et al., 2016). Alternatively, it is possible that the interaction occurs only when NTI needs to be moderated for the NTI signaling pathway to return to a resting state.

The molecular mechanisms underlying CNL- and TNL-mediated NTI activation are still not well understood (Chia & Carella, 2023). Recent reports indicate that certain NLRs have the ability to self-associate in response to effector protein binding, generating a ‘resistosome’, a multimeric complex that promotes NTI (Huang et al., 2023). Both CNLs and TNLs can form resistosome-like structures (Ma et al., 2020; Martin et al., 2020; Wang et al., 2019). While CNLs function as membrane-localized channels associated with calcium (Ca^2+^) influx and cell death, TNL resistosomes have NADase enzymatic activities that promote cell death (Huang et al., 2023). Our co-expression of Pic14 with cell death inducers in *N. benthamiana* suppressed cell death mediated by CNL proteins Ptr1 and Prf but not by the TNL Roq1. One explanation for the activity of Pic14 as a suppressor of only CNL-induced MAPK activation and cell death is the role of Ca^2+^ intake in CNL-mediated NTI (Bi et al., 2021). For example, it is possible that calcineurin B-like interacting protein kinases (CIPKs) or Ca^2+^-dependent protein kinases (CDPKs) are coregulated with MAPKs and are affected by Pic14 during CNL-mediated NTI (Boudsocq et al., 2010; de la Torre et al., 2013; Romeis & Herde, 2014; Thulasi Devendrakumar et al., 2018). Although both CNL-and TNL-associated signaling triggers cell death through MAPK cascades, these cascades differ in their components and modes of operation, pointing to distinct regulatory mechanisms (Ngou et al., 2021; Su et al., 2018). NLR-mediated immune activation requires involvement of another branch of plant immunity termed pattern triggered immunity (PTI), where conserved moieties of pathogens are recognized by extracellular receptors to trigger a separate set of defense responses (Yuan et al., 2021). TNL activation boosts transcript and protein levels of PTI components, suggesting a mutual amplification mechanism (Ngou et al., 2021; Yuan et al., 2021). This so far does not appear to be the case with CNLs, where MAPK activation is independent of PTI activation (Ngou et al., 2021). Thus, divergent regulatory mechanisms between CNLs and TNLs might explain the relevance of Pic14 to CNL and not TNL-associated pathways.

Collectively, our results indicate that Pic14 negatively regulates NTI by interference with MAPK signaling and they support the central importance of MAPK signaling in the Pto/Prf response, and possibly Ptr1 (del Pozo et al., 2004; Ekengren et al., 2003; Oh et al., 2010; Oh & Martin, 2011b, 2011a; Pedley & Martin, 2003, 2004; Roberts et al., 2019; Sheikh et al., 2023; Zhang et al., 2024). There are only two other reports of PP2Cs regulating MAPK phosphorylation in an NTI context, with both related to AvrRpt2/RPS2-mediated MAPK signaling (Mine et al., 2017; Wang et al., 2023). We propose that this type of NTI regulation is not unique to AvrRpt2/RPS2 and Pto/Prf signaling but instead is likely to be common and understudied. This hypothesis is supported by RNA-seq data analysis by Sobol et al. (2022), that showed that transcript abundance of 25 of the 97 tomato PP2Cs were altered during the NTI response. In conclusion, the present work expands the current model for Pto/Prf-induced signaling to include an additional negative regulatory component that monitors phosphorylation status of proteins in a MAPK cascade.

## EXPERIMENTAL PROCEDURES

### RT-qPCR

Five-week-old tomato plants of the indicated genotypes were vacuum infiltrated with *Pseudomonas syringae* pv. *tomato* (*Pst*) DC3000 or *Pst* DC3000Δ*avrPto*Δ*avrPtoB* suspensions of 5×10^6^ colony-forming units (cfu) mL^−1^. After 6 hr, 100 mg of leaf tissue was collected, and total RNA was extracted using the TRIzol reagent (Invitrogen). One μg of total RNA was used for cDNA synthesis with oligo(dT) primers using the RevertAid first strand cDNA synthesis kit (Thermo Fisher Scientific). RT-qPCR was performed with gene-specific primers (Supplemental Table S2) and the Fast SYBR green master mix (Applied Biosystems) using the QuantStudio 1 real-time PCR system (Applied Biosystems). The cycling conditions were set to 50°C for 2 min, 95°C for 10 min, and 40 cycles of 95°C for 30 sec, 56°C for 30 sec, and 72°C for 30 sec. Gene expression was determined based on the comparative Ct method (Pfaffl, 2001) using the QuantStudio Design & Analysis Software v1.5.2.

### Generation of tomato RG-pic14 mutants using CRISPR/Cas9

Tomato mutants were generated in Rio Grande-PtoR which expresses Pto and Prf which mediate recognition of AvrPto and AvrPtoB (Pedley & Martin, 2003a). For CRISPR/Cas9 mutagenesis, a guide RNA (gRNA) sequence (GCTCTAATAAGGTTACTCGG) targeting the exon 2 of *Pic14* was designed using the CRISPR-P website (http://crispr.hzau.edu.cn/CRISPR2/; Liu et al., 2017). The gRNA cassette was cloned into the p201N:Cas9 binary vector (Jacobs et al., 2017), and transformed into *Agrobacterium tumefaciens* strain AGL1. Plant transformation was performed at The Boyce Thompson Institute for Plant Research transformation facility. Mutations in the transformed plants were identified by amplifying genomic regions flanking the gRNA-targeted sites using specific primers (Table S2) and Sanger sequencing. Sequenced DNA was aligned against the wild-type *Pic14* sequence using the Geneious R11 software (Kearse et al., 2012).

### *Pseudomonas* inoculation assay

*Pst* DC3000Δ*avrPtoB* suspensions in 10 mM MgCl_2_ and 0.008% (v/v) Silwet L-77 were vacuum-infiltrated into four-week-old tomato plants of the indicated genotypes to assess bacterial growth, disease progression, and appearance of the hypersensitive response (HR) in leaves. For bacterial growth and tracking of disease progression, DC3000Δ*avrPtoB* was inoculated at 2×10^5^ cfu mL^−1^. Bacterial populations were measured at 3 hr (0 days post-inoculation; dpi) and at 2 dpi. Photographs of disease symptoms were taken 6 days after bacterial inoculation.

### *Agrobacterium*-mediated transient protein expression

*Agrobacterium tumefaciens* strains GV2260 carrying the indicated vectors were grown overnight in Lysogeny broth (LB) medium at 28°C. Bacteria were centrifuged at 4,000 rpm for 20 min and resuspended in induction medium (10 mM MgCl_2_ and 10 mM MES buffer [pH 5.6]). The OD_600_ of the resuspended cultures was adjusted to 0.4 unless stated otherwise and incubated in a shaker for 4 hr at 22°C with 200 μM acetosyringone. The bacterial suspension was then infiltrated into the abaxial surface of *Nicotiana benthamiana* or tomato leaves. To induce estradiol-mediated protein expression in *N. benthamiana*, leaf abaxial surfaces were sprayed with a 5 μM β-estradiol and 1% Tween 20 (v/v) solution. In tomato, estradiol-mediated protein expression was achieved by dipping whole leaves in the same solution for five minutes.

### Plant protein extraction

Total protein extraction from *N. benthamiana* or tomato leaves was carried out two days after agroinfiltration using a protein extraction buffer containing 50 mM Tris-HCl (pH 7.8), 200 mM NaCl, 1 mM EDTA, 2 mM DTT, 10% glycerol, and 1 mM phenylmethylsulfonyl fluoride (PMSF). Subsequently, 0.1% (v/v) Triton X-100 was added to each homogenized sample for a 10 min incubation on ice. Samples were centrifuged at 14,000 rpm for 30 min at 4°C. The supernatant was mixed with 3X Laemmli sample buffer and incubated at 95^0^C for 5 min.

### Immunoblotting

Proteins were resolved on a 10% sodium dodecyl-sulfate polyacrylamide gel electrophoresis (SDS-PAGE) gel and blotted onto polyvinylidene fluoride (PVDF) membrane using the Trans-blot turbo transfer system (Bio-Rad). PVDF membranes were incubated with anti-HA (1:5,000; v/v; Sigma-Aldrich) or anti-Luciferase (1:4,000; v/v; Sigma-Aldrich) primary antibodies and HRP conjugated anti-mouse (1:10,000; v/v; Jackson ImmunoResearch) or anti-rabbit (1:10,000; v/v; Jackson ImmunoResearch) secondary antibodies. Proteins were detected using the Clarity max ECL substrate (Bio-Rad) with the ImageQuant 800 imaging system (Amersham).

### Cell death and ion leakage measurement

To monitor plant cell death, 5-week-old *N. benthamiana* or tomato leaves were agroinfiltrated with bacterial strains carrying constructs of YFP, Pic14, or the Pic14AQ/NN variants at an OD_600_ of 0.4. The next day, the indicated cell death-triggering elicitors AvrPtoB_1-387_, Pto^Y207D^, AvrRpt2, and M3Kα^KD^ were infiltrated into the same infiltration areas at OD_600_ of 0.2, except for Mkk2 which was infiltrated in tomato leaves at OD_600_ of 0.4. Leaf images were captured at 3 to 7 days post agroinfiltration of the cell death-triggering elicitors using a Nikon D7100 camera equipped with an AF-S Micro NIKKOR 85mm 1:3.5G ED lens. XopQ-triggered cell death was monitored by measuring fluorescence of a co-expressed GFP protein (Schultink et al., 2017b). GFP fluorescence was visualized using the Fusion SOLO X imaging system (Vilber Lourmat). For ion leakage measurement experiments, leaf discs were collected at the indicated time points after elicitor treatments and floated in 10 mL tubes containing 5 mL of double-distilled water for 4 hours with constant shaking at 25°C. Ion leakage was measured with a DDS-12DW conductivity meter (BANTE Instruments, Shanghai, China).

### MAPK activity assay

*Agrobacterium tumefaciens* strain GV2260 carrying vectors for expression of HA-tagged YFP, Pic14, or the Pic14AQ/NN variants were infiltrated into *N. benthamiana* or tomato leaves. The following day, leaves were again agroinfiltrated with bacteria carrying pER8 vectors driving the estradiol-inducible expression of M3Kα^KD^, Mkk2, or AvrPto in the same infiltrated areas at an OD_600_ of 0.2 or 0.4 in *N. benthamiana* or tomato leaves, respectively. Ninety mg of leaf tissue was collected for total protein extraction in 100 mL of extraction buffer containing 50 mM Tris-HCl (pH 7.8), 200 mM NaCl, 1 mM EDTA, 10 mM NaF, 1 mM Na_2_MoO_4_, 2 mM Na_3_VO_4_, 10% glycerol, 2 mM DTT, and 1 mM PMSF. Protein samples were separated using a 10% acrylamide SDS-PAGE gel and immunoblotted with anti-phospho-p44/42 MAPK (Erk1/2) (anti-pMAPK; Cell Signaling Technology).

### Split luciferase complementation assay

Full-length gene fragments encoding Mkk1, Mkk2, Mpk1, Mpk2 and Mpk3 or Pic14 and its AQ/NN variants were PCR amplified using gene-specific primers (Table S2) and cloned in-frame to firefly luciferase fragments in the pCAMBIA1300:N-LUC or pCAMBIA1300:C-LUC binary vectors, respectively (Chen et al., 2008). The cloned plasmids were transformed into *Agrobacterium tumefaciens* strain GV2260 and co-infiltrated in *N. benthamiana* leaves. Following this, three mm-diameter leaf discs were punched from the infiltrated areas at 48 hr after agroinfiltration and floated overnight in 100 µL of double-distilled water in a white 96-well plate (Thermo Fisher Scientific). The next day, water was replaced with 100 µL of a 1 mM D-luciferin (Sigma-Aldrich) solution and incubated in the dark for 10 min. Luminescence was measured using a Veritas Microplate Luminometer (Promega). Whole leaf luminescence images of the infiltrated *N. benthamiana* leaves were captured as described by Sobol et al. (2022).

### Co-immunoprecipitation

*Agrobacterium tumefaciens* strain GV2260 carrying nLUC tagged Mkk1 or Mkk2 and HA-tagged YFP, Pic14, or the Pic14AQ/NN variants were co-infiltrated into *N. benthamiana* leaves. After 48 hr, 600 mg of leaf tissue was collected for each sample and used for protein extraction by adding 1.2 mL of extraction buffer containing 50 mM Tris-HCl (pH 7.8), 150 mM NaCl, 10% glycerol (v/v), 2 mM DTT, 0.1% NP-40 (v/v), 1 × protease inhibitor cocktail (Sigma-Aldrich), and 1 mM PMSF. Next, samples were treated with 0.1% Triton X-100 (v/v) and incubated for five min at 4 °C. Samples were then centrifuged at 4 °C at 14,000 rpm for 30 min. The supernatants (input) were collected at this stage and kept on ice. Next, 100 μL of HA-tag antibody-agarose beads (Sigma-Aldrich) pre-equilibrated in extraction buffer or 5 μL of anti-LUC antibodies (Sigma-Aldrich) were added to 1 mL of the supernatant. Protein immunoprecipitation using anti-HA beads was carried out for 6 hr at 4°C with gentle end-over-end rotation. For immunoprecipitation using anti-LUC antibodies, the samples were further incubated with Protein A agarose beads (Thermo Fisher Scientific) pre-equilibrated in extraction buffer for 16 hr at 4°C. The beads were collected by centrifugation at 14,000 rpm for 10 min at 4°C. The beads were washed three times in extraction buffer and twice in extraction buffer with 0.5% NP-40 (v/v). Last, the beads were incubated in 1X Laemmli sample buffer at 70°C for 5 min.

### Purification of GST-tagged fusion proteins

*Pic14* and its AQ/NN variants were cloned between the SalI and NotI sites in the pGEX-4T1 plasmid (Amersham) and mobilized into the Rosetta *E. coli* strain (Novagen). Transformed *E. coli* cells expressing GST, GST-Pic14, GST-Pic14AQ, and GST-Pic14NN were grown in 2x Yeast extract tryptone (2x YT) medium overnight at 37°C. The next day, bacterial cultures were inoculated in 250 mL of 2x YT media and grown at 37°C to an OD_600_ between 0.6 and 0.8. Following this, 0.5 mM isopropyl ß-D-1-thiogalactopyranoside (IPTG) was added to induce protein expression for 6 hr at 28°C. Bacterial cells were pelleted by centrifugation at 4,000 rpm for 20 min at 4°C. Bacterial pellets were then suspended in lysis buffer containing 25 mM Tris-HCl (pH 7.5), 150 mM NaCl, 5 mM MgCl_2_, 1 mM PMSF, 0.1% Triton X-100 (v/v), 1 mg/mL lysozyme, and 20 μg/mL DNase I and subjected to end-over-end rotation for 30 min at 4°C. The lysed bacterial suspension was centrifuged for 30 min at 4,000 rpm at 4 °C. The supernatant was passed through a column containing glutathione resin (GenScript) pre-washed in GST binding buffer (25 mM Tris-HCl [pH-7.5], 150 mM NaCl, and 5 mM MgCl_2_). The beads were washed three times with a 25 mM Tris-HCl (pH 7.5) and 150 mM NaCl solution. Finally, the bound proteins were eluted using an elution buffer containing 50 mM Tris-HCl (pH 8.0), 5% glycerol, and 20 mM reduced glutathione.

### Phosphatase activity assay

*In vitro* phosphatase activity of GST, GST-Pic14, and GST-Pic14AQ/NN variants was determined using a Ser/Thr phosphatase assay system (V2460; Promega) according to the manufacturer’s instructions. Five hundred ng of recombinant GST-fusion proteins were incubated with 5 μL of 1 mM phosphopeptide [RRA(phosphoT)VA] and 10 μL of 5×PP2C reaction buffer (250 mM imidazole, pH 7.2, 1 mM EGTA, 25 mM MgCl_2_, 0.1% β-mercaptoethanol (v/v), and 0.5 mg/mL BSA) at 25°C for 15 min in a transparent 96-well plate. The reaction was stopped by adding 50 μL of a molybdate dye/additive mixture supplied with the kit. For substrate dephosphorylation assays, 250 ng of purified GST or GST-Mkk2 proteins were incubated in a kinase activity promoting buffer containing 25 mM Tris-HCl (pH 7.5), 2 mM dithiothreitol (DTT), and 10 mM MgCl_2_ in the presence or absence of 100 µM ATP for 30 min at 30°C. The reaction mixtures were incubated with 500 ng of purified GST, GST-Pic14, or its variants in a 5×PP2C reaction buffer for another 15 min at 22°C. Finally, 50 μL of molybdate dye/additive mixture was added to each reaction to stop it. The absorbance of the reaction mixture was measured at 630 nm in a microplate reader (Synergy HT, BioTek).

### Bioinformatic analyses

Significant amino acids within the KIM and PP2C domains were identified using a multiple sequence alignment of Pic14 and its homologs performed using ClustalW (https://www.genome.jp/tools-bin/clustalw) with default parameters. Phylogenetic trees were generated using the MEGA X software (Kumar et al., 2018) according to the neighbor joining method (bootstrap value set to 1,000 iterations) with the parameters of the JTT matrix-based model (Jones et al., 1992).

## ACKNOWLEDGEMENTS

We thank Joyce Van Eck, Qingzhen Jiang, and Tish Keen for tomato transformation; Tara McCrudden, Julie Bell, Kelly Jackson, Philip Ricci, and Nick Vail for plant care. JC thanks Professor Amir Sharon for supporting his research at Tel Aviv University. Research funding was provided by grant IS-5362-21CR to GSe and GBM from the United States-Israel Binational Agricultural Research and Development Fund (BARD). This paper is dedicated to the memory of our beloved Professor Guido Sessa.

## AUTHOR CONTRIBUTIONS

GSe, JC, GSo, and GBM conceived and designed the experiments. FX and NZ designed gRNAs and constructed vectors and oversaw generation of the tomato mutants. JC performed phenotyping, molecular, and biochemical experiments. FX helped with genotyping and phenotyping experiments in tomato. GSe, JC, GSo, FX, NZ, and GBM interpreted the data. All authors, except GSe, contributed to the writing, and read and approved the manuscript.

## SUPPORTING INFORMATION

**Figure S1.** Expected truncated variants of Pic14 produced in RG-pic14 mutants.

**Figure S2.** Amino acids essential for interaction with MAPKs and phosphatase activity are conserved between Pic14 and Arabidopsis PP2Cs.

**Figure S3.** Detection of *Agrobacterium*-mediated transient expression of 4X-hemagglutinin (HA)-tagged YFP, Pic14, Pic14AQ, and Pic14NN variants in *Nicotiana benthamiana* leaves by immunoblotting.

**Figure S4.** Pic14 does not suppress XopQ-induced cell death in *N. benthamiana*.

**Figure S5.** Protein expression of HA-tagged M3Kα^KD^, Mkk2, YFP, Pic14, and its variants in *N. benthamiana* and tomato leaves by immunoblotting.

**Figure S6.** Phylogeny of Pic14 homologs and their interactors.

**Figure S7.** Western blot for detection of proteins used in the split-luciferase assay.

**Figure S8.** *In vitro* phosphatase assay for Pic14 variants.

