## Supplemental Information for "PP2C phosphatase Pic14 negatively regulates tomato Pto/Prf-triggered immunity by inhibiting MAPK activation"

**Table S1.** Differential expression of the *Pic14* gene during the Pto/Prf NTI response.

| | **RG-PtoR / RG-prf3** | **pFDR** | **RG-PtoR / RG-prf19** | **pFDR** | **RG-PtoR / RG-prf3** | **pFDR** | **RG-PtoR / RG-prf19** | **pFDR** |
|---|---|---|---|---|---|---|---|---|
| **Timepoint** | **4 h** | | **4 h** | | **6 h** | | **6 h** | |
| **Fold-change / pFDR** | 3.15 | 3.09E-10 | 3.46 | 3.34E-11 | 3.82 | 7.86E-14 | 6.63 | 2.41E-16 |

RG-PtoR has a functional Pto/Prf pathway. RG-prf3 and RG-prf19 have a mutation in Prf which abolishes Pto/Prf NTI.

Plants were inoculated with Pst DC3000 which activates NTI when Pto/Prf are present.

Data are summarized in Sobol et al., MPMI (2022). 35:737-747. Supplemental Table S5.

Data are from Pombo et al., 2014 DOI: 10.1186/s13059-014-0492-1, data extracted from the Tomato Functional Genomics Database website: http://ted.bti.cornell.edu/

**Table S2.** Primers used in this study.

| Primer name | Gene Solyc ID | Primer sequence (5'-3') | Purpose |
|---|---|---|---|
| Pic14_F | Solyc06g082080 | CGGCTTCTCTTCGTTGTAAAG | Genotyping CRISPR-generated mutants |
| Pic14_R | Solyc06g082080 | CTTGAGGGATTACACCGGTT | |
| Pic14-qRT_F | Solyc06g082080 | CCGGCTTCTCTTCGTTGTAA | RT-qPCR |
| Pic14-qRT_R | Solyc06g082080 | TCTGAAGCCGAAGAATCCGT | |
| UDPG1-qRT_F | Solyc09g092500 | TTGGACAGATCAAGGGACTAATG | |
| UDPG1-qRT_R | Solyc09g092500 | CACTCTCAACCACACCATCTT | |
| UDPG2-qRT_F | Solyc10g085880 | CCTGGATTGTTCGACAAGAT | |
| UDPG2-qRT_R | Solyc10g085880 | CTCCTCCGCTTTCTTCATTT | |
| Laccase-qRT_F | Solyc04g072280 | AACGTCCCGATCGTAGAA | |
| Laccase-qRT_R | Solyc04g072280 | GGATGATCAACTCCACCTAATAA | |
| SlEF1α-qRT_F | Solyc06g005060 | TCCAAAGATGGTCAGACCCGTGAA | |
| SlEF1α-qRT_R | Solyc06g005060 | ATACCTAGCCTTGGAGTACTTGGG | |
| 2XHA-SalI_F | | AAGTCGACaTATCCATATGATGTTCCAGATTATGCTTATCCATATGATGTTCCAGATTATGCTGACGTCAA | Expression of HA-tagged proteins in plant cells |
| 2XHA-PstI_R | | TTGACGTCAGCATAATCTGGAACATCATATGGATAAGCATAATCTGGAACATCATATGGATAtGTCGACTT | |
| HA-PstI_F | | TAAACTGCAGTATCCATATGATGTTCCAGATTATGCTTGAGATCGTTCAAACATTTGGCAATAAAGTTTC | |
| NosT-HindII_R | | AAAAAGCTTGATCTAGTAACATAGATGACAC | |
| YFP-KpnI_F | | ATTAAGGTACCATGGTGAGCAAGGGCGAG | |
| YFP-HA-XbaI_R | | ATTATTCTAGAAGCGTAATCTGGAACATCGTATGGGTACTTGTACAGCTCGTCCATGCC | |
| Pic14-KpnI_F | Solyc06g082080 | AACGGGGTACCCATCAGAAATCGAAAAAA | |
| Pic14-HA-SalI_R | Solyc06g082080 | ATTATGTCGACAGCGTAATCTGGAACATCGTATGGGTAACAGAATCGTCCCAGTTG | |
| Pic14-AQ_F | Solyc06g082080 | TCTACGGTTTTGGCGCAGAAGAGACCGACT | Site-directed mutagenesis |
| Pic14-AQ_R | Solyc06g082080 | AAACCGTAGAAGTAGAACAATCAGTATTAG | |
| Pic14-D160N_F | Solyc06g082080 | TTTTTTGGTATATTTAATGGTCATGGAGGAGTG | |
| Pic14-D160N_R | Solyc06g082080 | TACCAAAAAAACCCTGTTTGGAATCTCCTT | |
| Pic14-D234N_F | Solyc06g082080 | TCCAATGCTGGGAATTGTCGTGCTGTTG | |
| Pic14-D234N_R | Solyc06g082080 | CAGCATTGGATACAACTAGATTGCCCTTT | |
| C-Luc-Pic14-KpnI_F | Solyc06g082080 | AACGGGGTACCCATCAGAAATCGAAAAAA | Split-Luciferase Complementation Assay |
| C-Luc-Pic14-SalI_R | Solyc06g082080 | ATTATGTCGACACAGAATCGTCCCAGTTG | |
| N-Luc-Mkk1-KpnI_F | Solyc12g009020 | ATTATGGTACCATGAAGAAAGGATCTTTT | |
| N-Luc-Mkk1-SalI_R | Solyc12g009020 | AACGCGTCGACTAGCTCAGTAAGTGTTGC | |
| N-Luc-Mkk2-SalI_F | Solyc03g123800 | ATTATGTCGACATGCGACCAGCCGCCAAC | |
| N-Luc-Mkk2-SalI_R | Solyc03g123800 | AACGCGTCGACAGAAGAGGAGGAAAAATGA | |
| N-Luc-Mpk1-SacI_F | Solyc12g019460 | TATATGAGCTCATGGATGGTTCCGTTCCGC | |
| N-Luc-Mpk1-BamHI_R | Solyc12g019460 | TTAGCGGATCCCATGCGCTGGTATTCAGGA | |
| N-Luc-Mpk2-SacI_F | Solyc08g014420 | ATTATGAGCTCATGGATGGTTCAGCTCCGC | |
| N-Luc-Mpk2-BamHI_R | Solyc08g014420 | AACGCGGATCCCATGTGCTGGTATTCGGGA | |
| N-Luc-Mpk3-KpnI_F | Solyc06g005170 | ATTATGGTACCATGGTTGATGCTAATATG | |
| N-Luc-Mpk3-SalI_R | Solyc06g005170 | AACGCGTCGACAGCATATTCAGGATTCAA | |
| Pic14-SalI_F | Solyc06g082080 | ATTATGTCGACTGTCTTGCACCGTCGCT | GST-tagged protein purification |
| Pic14-NotI_R | Solyc06g082080 | AACGCGCGGCCGCACAGAATCGTCCCAGTT | |

**RG-PtoR**

MSCTVALSNSPVFSPSRVPVPASLRCKVSSSSSSSSPETLNLTH
SPSKTSSSPSSPSSPLRILRLQKPPPSNLIRASNTDCSTSTVLKR
KRPTRLDLPVASMSFGNFPVTPAGVADLVEVEGDGYSVCCKRG
RKGAMEDRHSAMVNLKGDSKQGFFGIFDGHGGVKAAEFSAEN
LNKNIMNELGKTTDDKIEVAVKNGYLKTDTEFLSQEVRGGSCCV
TALIQKGNLVVSNAGDCRAVVSRGGLAEALTSDHRPSRKDEKDR
IEASGGYVDCCHGVWRIQGSLAVSRGIGDQYLKQWVTAEPETKI
LELNPELEFLVLASDGLWDTVSNQEAVDIARPLCTGISTQQPLSA
CRKLIDLSVSRGSLDDISVLIIQLGRFC*

**RG-pic14-1**

MSCTVALSNSPVFSPSRVPVPASLRCKVSSSSSSSSPETLNLTH
SPSKTSSSPSSPSSPLRILRLQKPPPVTLLELLILIVLLLRF*

**RG-pic14-2**

MSCTVALSNSPVFSPSRVPVPASLRCKVSSSSSSSSPETLNLTH
SPSKTSSSPSSPSSPLRILRLQKPPP*

**Figure S1. Expected truncated variants of Pic14 produced in RG-pic14 mutants.** Amino acid sequences of Pic14 and the possible variants produced in wild-type (RG-PtoR) and the RG-pic14-1 and RG-pic14-2 mutants due to an early stop codon. The RG-pic14 lines contain a premature stop codon at the 87th and 71st amino acids of the Pic14 protein. Asterisks (*) denote the stop codon.

```
AtPP2C5         MQLSKNPIKQTRNREKNYTDDFTMKRSVIMAPESPVFFPPPLV----------------- 43
Pic14           -----------------------MSC-TVALSNSPVFSPSRVPVPASLRCKVSSSSSSSS 36
AtAP2C1         -----------------------MSC-SVAVCNSPVFSPSSSL-----FCNKSSILS-SP 30
                                       *.   :   :**** *

AtPP2C5         ---------------FSPTSVKTPLSSPRSSPPKLTMVACPP-------RKPKETKTTGS 81
Pic14           PETLN--LTH------SPSKTSSSPSSPSSPLRILRLQKPPPSNLI-------RASNTDC 81
AtAP2C1         QESLSLTLSHRKPQTSSPSSPSTTVSSPKSPFR-LRFQKPPSGFAPGPLSFGSESVSASS 89
                                **:. .:  *** *    * :   *            .: .:..
                          KIM                                 PP2C domain
AtPP2C5         DSETVIKRKRPPMLDLTAAPTVASW--C--STTRETAEKGAEVVEAEEDGYYSVYCKRGR 137
Pic14           STSTVIKRKRPTRLDLPVASMSFGN-------FPVTPAGVADLVEVEGD-GYSVCCKRGR 133
AtAP2C1         PPGGVIKRKRPTRLDIPIGVAGFVAPISSSAAVAATPREECREVEREGD-GYSVYCKRGR 148
                    *******  **:  .                *    .  ** * *  *** *****
                                           PP2C domain
AtPP2C5         RGPMEDRYFAAVDRNDDGGYKNAFFGVFDGHGGSKAAEFAAMNLGNNIEAAMASARSGED 197
Pic14           KGAMEDRHSAMVN--LKGDSKQGFFGIFDGHGGVKAAEFSAENLNKNIMNEL-GKT---T 187
AtAP2C1         REAMEDRFSAITN--LHGDRKQAIFGVYDGHGGVKAAEFAAKNLDKNIVEEVVGKR---D 203
                :  ****. * .:   .*. *:.:**::***** *****:* **.:**   : .
                                           PP2C domain
AtPP2C5         GCSMESAIREGYIKTDEDFLKE-GSRGGACCVTALISKGELAVSNAGDCRAVMSRGGTAE 256
Pic14           DDKIEVAVKNGYLKTDTEFLSQ-EVRGGSCCVTALIQKGNLVVSNAGDCRAVVSRGGLAE 246
AtAP2C1         ESEIAEAVKHGYLATDASFLKEEDVKGGSCCVTALVNEGNLVVSNAGDCRAVMSVGGVAK 263
                  .:  *::.**: ** .**.:   :**:******:.:*:*.**********:* ** *:
                                           PP2C domain
AtPP2C5         ALTSDHNPSQANELKRIEALGGYVDCCNGVWRIQGTLAVSRGIGDRYLKEWVIAEPETRT 316
Pic14           ALTSDHRPSRKDEKDRIEASGGYVDCCHGVWRIQGSLAVSRGIGDQYLKQWVTAEPETKI 306
AtAP2C1         ALSSDHRPSRDDERKRIETTGGYVDTFHGVWRIQGSLAVSRGIGDAQLKKWVIAEPETKI 323
                **:***.**: :* .***: *****  :*******:*********  **:** *****:
                                           PP2C domain
AtPP2C5         LRIKPEFEFLILASDGLWDKVTNQEAVDVVRPYCVGVENPMTLSACKKLAELSVKRGSLD 376
Pic14           LELNPELEFLVLASDGLWDTVSNQEAVDIARPLCTGISTQQPLSACRKLIDLSVSRGSLD 366
AtAP2C1         SRIEHDHEFLILASDGLWDKVSNQEAVDIARPLCLGTEKPLLLAACKKLVDLSASRGSSD 383
                 .:: : ***:********.*:******:.** * * ..   *:**:** :**..*** *
                PP2C domain
AtPP2C5         DISLIIIQLQNFLP 390
Pic14           DISVLIIQLGRFC- 379
AtAP2C1         DISVMLIPLRQFI- 396
                ***:::* * .*
```

**Figure S2. Amino acids essential for interaction with MAPKs and phosphatase activity are conserved between Pic14 and Arabidopsis PP2Cs.** Multiple sequence alignments between Pic14 and Arabidopsis PP2Cs showing functional kinase-interacting motif (KIM) and PP2C domains. Identification of the KIM was based on the consensus sequence $[K/R]_{(3-4)}$-$X_{(1-6)}$-[L/I]-X-[L/I] (Brock et al., 2010; Schweighofer et al., 2007). The SMART tool (http://smart.embl-heidelberg.de/) was used to delineate the PP2C domain. Conserved amino acids within the KIM and PP2C domains between Pic14 and Arabidopsis PP2Cs are highlighted in yellow. Red and green boxes indicate amino acid sequences that make up the KIM and PP2C domains, respectively.

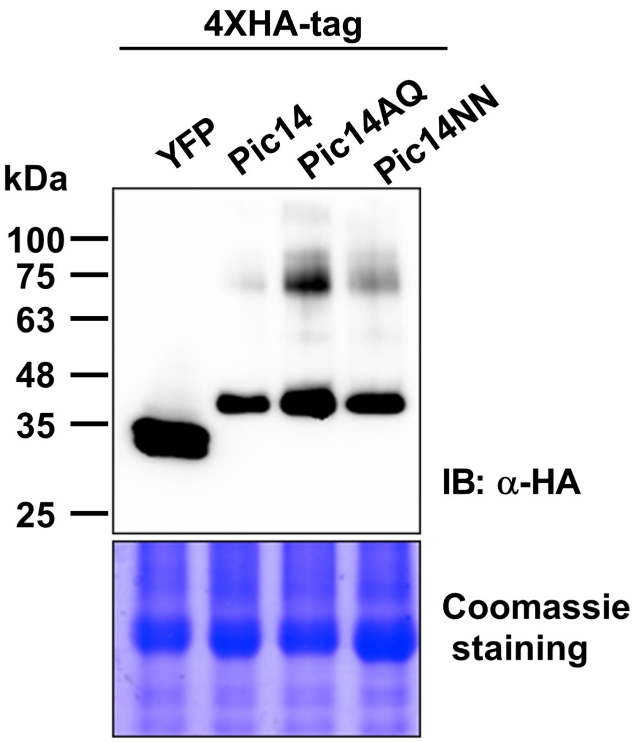


**Figure S3. Detection of *Agrobacterium*-mediated transient expression of 4X-hemagglutinin (HA)-tagged YFP, Pic14, Pic14AQ, and Pic14NN variants in *Nicotiana benthamiana* leaves by immunoblotting.** *Agrobacterium tumefaciens* GV2260 strains containing constructs of HA-tagged YFP, Pic14, Pic14AQ, and Pic14NN variants were syringe-infiltrated at $OD_{600}$ = 0.4 into leaves of five-week-old *N. benthamiana* plants. Two days later, the infiltrated leaves were used for total protein extraction and immunoblot analysis. The upper panel shows the protein bands detected by immunoblotting with an anti-HA antibody. The lower panel shows coomassie-stained gel as a loading control.

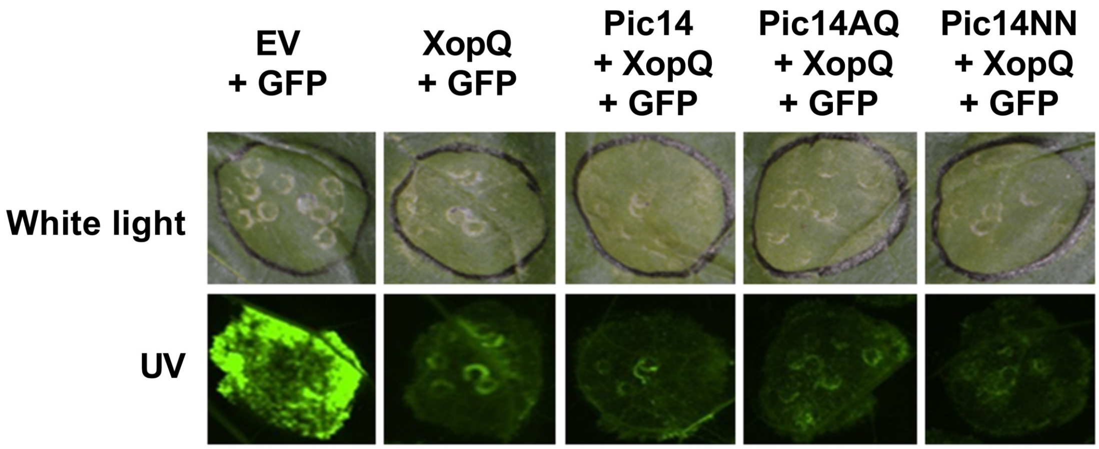


**Figure S4. Pic14 does not suppress XopQ-induced cell death in *N. benthamiana*.** *A. tumefaciens* GV2260 strains carrying plasmids of empty vector (EV), Pic14, Pic14AQ, or Pic14NN variants were syringe-infiltrated ($OD_{600}$ = 0.4) into leaves of five-week-old *N. benthamiana* plants. The next day, co-infiltration was carried out with GV2260 strains harboring constructs of green fluorescent protein (GFP) and *Xanthomonas* effector Xanthomonas outer protein Q (XopQ) at $OD_{600}$ = 0.2 in the same areas of *N. benthamiana* leaves. The agroinfiltrated *N. benthamiana* leaves were imaged 3 days post-infiltration under white light (upper panel) and UV light (lower panel). The images shown here are representative of six technical replicates from two biological experiments.

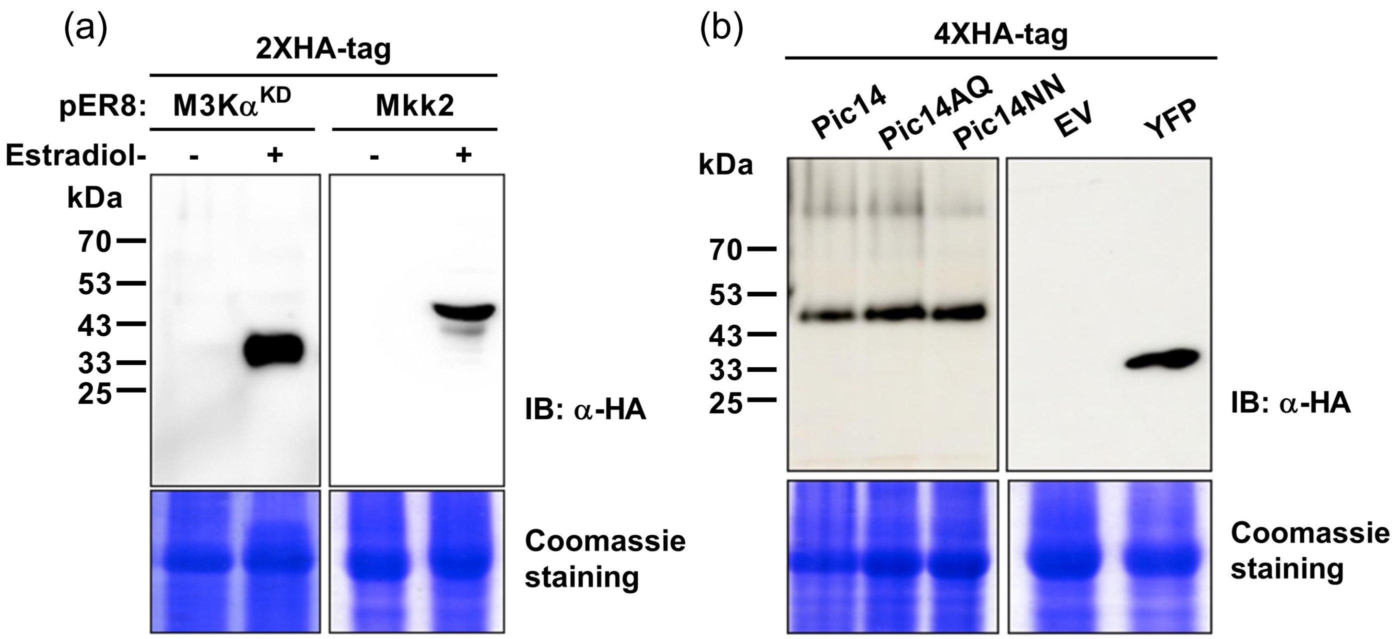


**Figure S5. Protein expression of HA-tagged M3Kα$^{KD}$, Mkk2, YFP, Pic14, and its variants in *N. benthamiana* and tomato leaves by immunoblotting.** (a) Estradiol-inducible expression of HA-tagged M3Kα$^{KD}$ and Mkk2 in *N. benthamiana* and tomato leaves. *A. tumefaciens* GV2260 strains containing constructs of pER8:M3Kα$^{KD}$ and pER8:Mkk2 were infiltrated into *N. benthamiana* and tomato leaves at $OD_{600}$ = 0.4. Protein expression was detected 8 h after 5 μM estradiol treatment. The minus (-) and plus (+) signs indicate mock and estradiol-treated samples, respectively. Immunoblotting was performed with an α-HA antibody. (b) *A. tumefaciens* GV2260 strains containing constructs of YFP or Pic14, Pic14AQ, and Pic14NN variants were infiltrated into tomato leaves at $OD_{600}$ = 0.4. Total proteins were extracted from tomato leaves 48 h after agroinfiltration. Transient expression of 4HA-tag YFP or Pic14, Pic14AQ, and Pic14NN variants in tomato leaves was detected by immunoblotting with an α-HA antibody. EV, empty vector. In (a) and (b), coomassie-stained gels show equal protein loading.

(a)

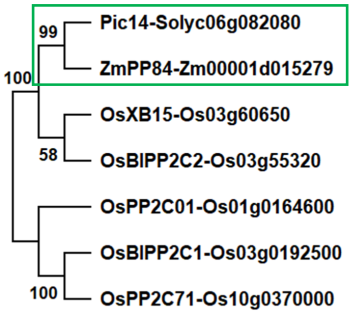


| Plants | Interactors | References |
|---|---|---|
| Tomato | | This study |
| Maize | Mkk1 | Guo et al., 2023 |
| Rice | XA21 | Park et al., 2008 |
| Rice | | Hu et al., 2009 |
| Rice | | Akimoto-Tomiyama et al., 2018 |
| Rice | | Akimoto-Tomiyama et al., 2018 |
| Rice | | Akimoto-Tomiyama et al., 2018 |

(b)

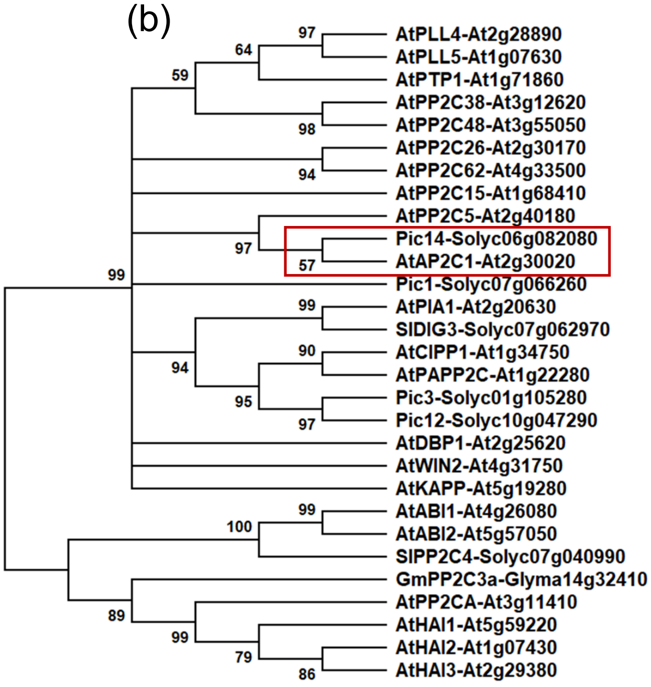


| Plants | Interactors | References |
|---|---|---|
| Arabidopsis | FLS2 and EFR | DeFalco et al., 2022 |
| Arabidopsis | FLS2 and EFR | DeFalco et al., 2022 |
| Arabidopsis | Mpk3 and Mpk6 | Bartels et al., 2009 |
| Arabidopsis | BIK1 | Couto et al., 2016 |
| Arabidopsis | | Couto et al., 2016 |
| Arabidopsis | | Akimoto-Tomiyama et al., 2018 |
| Arabidopsis | | Akimoto-Tomiyama et al., 2018 |
| Arabidopsis | BAK1 | Diao et al., 2024 |
| Arabidopsis | Mpk3, Mpk4, and Mpk6 | Brock et al., 2010 |
| Tomato | | This study |
| Arabidopsis | Mpk6 and Mpk4 | Schweighofer et al., 2007 |
| Tomato | Pti1b | Giska and Martin, 2019 |
| Arabidopsis | | Widjaja et al., 2009 |
| Tomato | | Tao et al., 2004 |
| Arabidopsis | CERK1 | Liu et al., 2018 |
| Arabidopsis | RPW8.2 | Wang et al., 2012 |
| Tomato | | Xia et al., 2024 |
| Tomato | | Xia et al., 2024 |
| Arabidopsis | Mpk11 | Carrasco et al., 2014 |
| Arabidopsis | | Lee et al., 2008 |
| Arabidopsis | FLS2 | Gómez-Gómez et al., 2001 |
| Arabidopsis | MAPKKK18 and CPK5 | Mitula et al., 2015; Yamada and Mine, 2024 |
| Arabidopsis | | Leung et al., 1997 |
| Tomato | SnRK2 and Sw-5b | Huang et al., 2024 |
| Soybean | | Seo et al., 2014 |
| Arabidopsis | | Lim et al., 2014 |
| Arabidopsis | Mpk3 and Mpk6 | Mine et al., 2017 |
| Arabidopsis | | Mine et al., 2017 |
| Arabidopsis | | Mine et al., 2017 |

**Figure S6. Phylogeny of Pic14 homologs and their interactors.** Phylogenetic tree of Pic14 and its homologs from (a) monocot and (b) dicot plants, alongside a column that indicates their protein interactors. The green and red boxes show Pic14 homologs from maize and Arabidopsis. Amino acid sequences of Pic14 and other PP2C phosphatases from rice, maize, soybean, tomato, and Arabidopsis were used to generate a neighbor-joining phylogenetic tree in MEGA X. MUSCLE program was used to build the sequence alignment. Bootstrap values were set to 1000 replicates. Branch lengths indicate the number of substitutions per site.

**Figure S7**

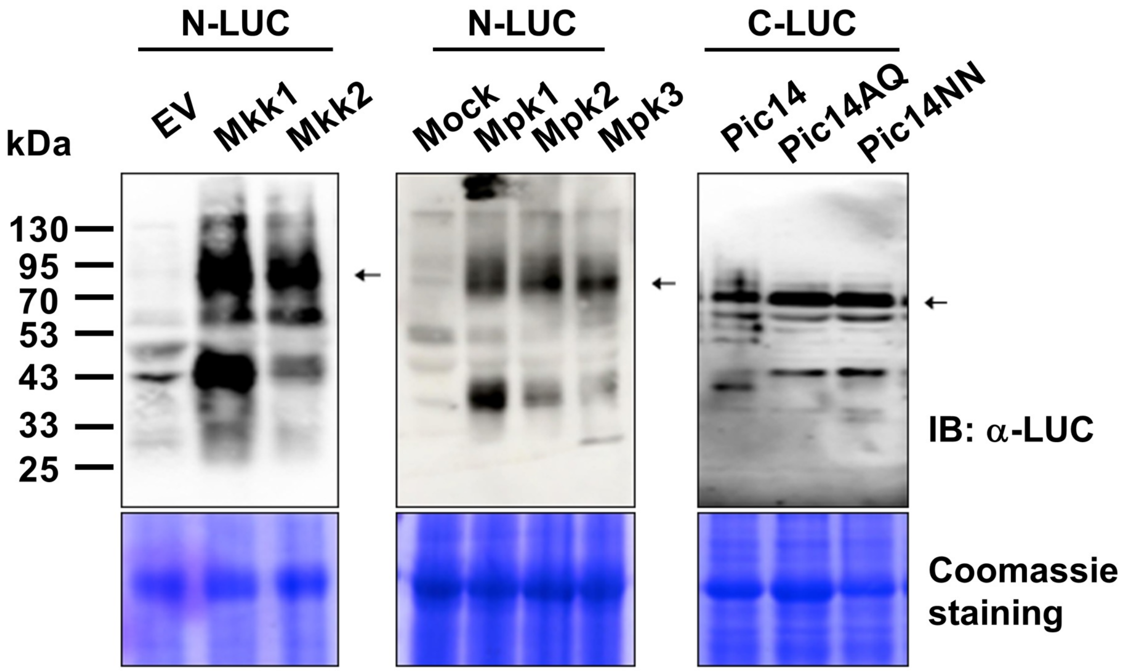


**Figure S7. Western blot for detection of proteins used in the split-luciferase assay.** *A. tumefaciens* GV2260 strains were used to express the Mkks, Mpks, or Pic14 and its variants fused to the N-terminal (N-LUC) or C-terminal (C-LUC) half of the luciferase protein in *N. benthamiana* leaves. 48 h after agroinfiltration, total proteins were extracted from the infiltrated areas of the *N. benthamiana* leaves. Protein expression was detected by immunoblot analysis with anti-luciferase (α-LUC) antibodies. Mock represents buffer-infiltrated leaves. Arrows indicate the specific protein bands.

**Figure S8. *In vitro* phosphatase assay for Pic14 variants.** (a) 10% Sodium dodecyl-sulfate polyacrylamide gel (SDS-PAGE) analysis of the purified glutathione S-transferase (GST) tagged Pic14, Pic14AQ, and Pic14NN variants or control GST proteins used for the phosphatase assay using coomassie staining. (b) 10% SDS-PAGE analysis of the purified GST-Mkk2 protein used for the phosphatase assay by coomassie staining. In (a) and (b), Arrows indicate the purified protein bands. (c) The flowchart shows the dephosphorylation assay of the GST-Mkk2 protein by GST-Pic14 using a phosphatase assay. Briefly, GST-Mkk2 protein was *in vitro* phosphorylated in a kinase assay buffer containing ATP and then incubated with control GST or GST-tagged Pic14, Pic14AQ, and Pic14NN variants. Next, the reaction was terminated by the addition of molybdate dye solution, and OD was measured at 630 nm. (d) Plate images showing dephosphorylation of GST-Mkk2 by GST-Pic14 in a phosphatase assay reaction. The green color of the solution indicates dephosphorylation of GST-Mkk2 by GST-Pic14, while the yellow color of the solution suggests no phosphatase activity. (e) Phosphatase-substrate assay with no ATP control. GST-Mkk2 protein was incubated in a kinase assay buffer without ATP and then used for phosphatase assay by control GST or GST-tagged Pic14, Pic14AQ, and Pic14NN variants. Bars represent means ± standard deviation (SD) of n = 3 reactions. ns, no significant difference by one-way ANOVA.

**Figure S8**

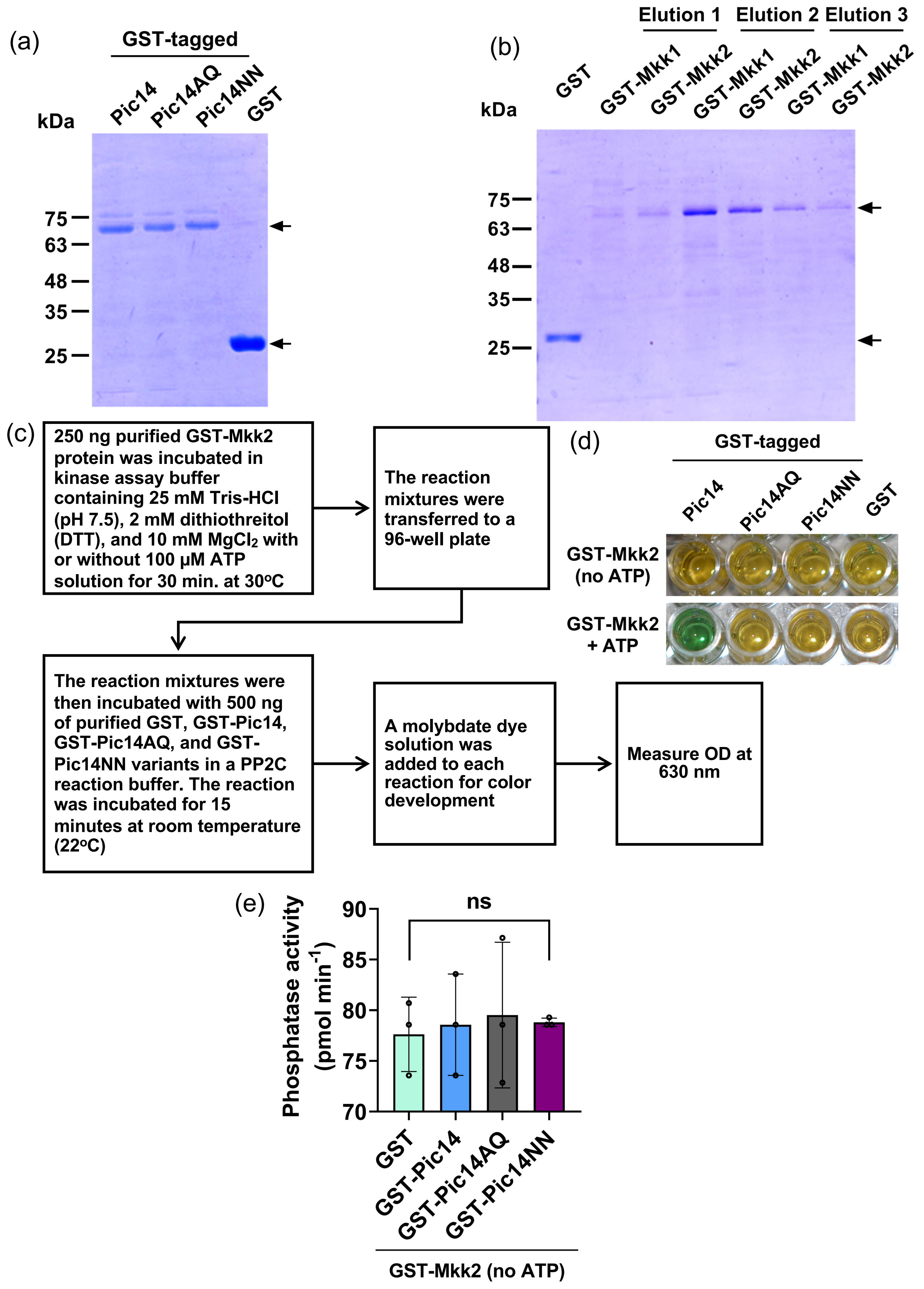

(a)
GST-tagged
kDa
Pic14
Pic14AQ
Pic14NN
GST
75
63
48
35
25
(b)
Elution 1
Elution 2
Elution 3
kDa
GST
GST-Mkk1
GST-Mkk2
GST-Mkk1
GST-Mkk2
GST-Mkk1
GST-Mkk2
75
63
48
35
25
(c)
250 ng purified GST-Mkk2 protein was incubated in kinase assay buffer containing 25 mM Tris-HCl (pH 7.5), 2 mM dithiothreitol (DTT), and 10 mM MgCl2 with or without 100 µM ATP solution for 30 min. at 30oC
The reaction mixtures were transferred to a 96-well plate
The reaction mixtures were then incubated with 500 ng of purified GST, GST-Pic14, GST-Pic14AQ, and GST-Pic14NN variants in a PP2C reaction buffer. The reaction was incubated for 15 minutes at room temperature (22oC)
A molybdate dye solution was added to each reaction for color development
Measure OD at 630 nm
(d)
GST-tagged
Pic14
Pic14AQ
Pic14NN
GST
GST-Mkk2 (no ATP)
GST-Mkk2 + ATP
(e)
ns
Phosphatase activity (pmol min-1)
90
85
80
75
70
GST
GST-Pic14
GST-Pic14AQ
GST-Pic14NN
GST-Mkk2 (no ATP)
